# Soil characteristics constrain the response of bacterial and fungal communities and hydrocarbon degradation genes to phenanthrene soil contamination and phytoremediation with poplars

**DOI:** 10.1101/2020.09.04.284042

**Authors:** Sara Correa-García, Karelle Rheault, Julien Tremblay, Armand Séguin, Etienne Yergeau

## Abstract

Rhizodegradation is a promising cleanup technology where microorganisms degrade soil contaminants in the rhizosphere. A symbiotic relationship is expected to occur between plant roots and soil microorganisms in contaminated soils that enhance natural microbial degradation in soils. However, little is known about how this initial microbiota influences the rhizodegradation outcome in a context of different soil microbiotas. Recent studies have hinted that soil initial diversity has a determining effect on the outcome of contaminant degradation. To test this hypothesis, we planted (P) or not (NP) balsam poplars (*Populus balsamifera*) in two soils of contrasting diversity (agricultural and forest) that were contaminated or not with 50 mg kg^-1^ of phenanthrene (PHE). The DNA from the rhizosphere of the P and the bulk soil of the NP pots was extracted and the bacterial genes encoding for the 16S rRNA, the PAH ring-hydroxylating dioxygenase alpha subunits (PAH-RHDα) of gram-positive (GP) and gram-negative (GN) bacteria, and the fungal ITS region were sequenced to characterize the microbial communities. and the abundance of the PAH-RHDα genes were also quantified by real-time quantitative PCR. Plant presence had a significant effect on PHE degradation only in the forest soil, whereas both NP and P agricultural soils degraded the same amount of PHE. Bacterial communities were principally affected by the soil type, and upon contamination the dominant PAH degrading community was similarly constrained by soil type. Our results highlight the crucial importance of soil microbial and physicochemical characteristics in the outcome of rhizoremediation.

## 1. Introduction

The expansion of the human population coupled with a burst in industrialization has led to a sensible increase in fossil fuel combustion (Jonsson 2012). As a result, there has been an increase in all types of pollutant emissions, including polycyclic aromatic hydrocarbons (PAH) (Jones et al. 1989; Ravindra, Sokhi et Van Grieken 2008). PAH are organic compounds formed by a variable number of fused aromatic rings that constitute a priority concern due to their adverse effects on environmental (Menzie, Potocki et Joseph 1992) and human health (Kim et al. 2013). PAH are ubiquitous in the environment and are often found polluting soils (Wilcke 2007). This is of special concern, since soil is one of the most important resources upon which many key human economic and social activities depend (Robinson et al. 2017).

To remediate soils, a wide range of clean up technologies have been developed, ranging from relatively simple but expensive excavation, transportation, washing and dumping of small affected soil areas (Kuppusamy et al. 2016) to more complex but comparatively cheaper phytoremediation strategies where the soil is cleaned up by the combined action of plant and microorganisms (Pilon-Smits 2005). Phytoremediation has proven to be effective to decontaminate soils polluted by PAH (Kuppusamy et al. 2017; Spriggs, Banks et Schwab 2005; Guo et al. 2018; Lu et Lu 2015). Among the many types of phytoremediation approaches, rhizodegradation is a genuine plant-microbe co-enterprise, with the potential to completely remove PAH from soils through the action of bacterial, fungal and plant enzymes in the rhizospheric environment (Correa-Garcia 2018).

Phenanthrene (PHE) is a low molecular weight PAH. It is formed by three fused rings and it is relatively easy to degrade by a wide range of microorganisms through aerobic or anaerobic metabolic pathways (Cerniglia 1993). Most of microbial degraders are bacteria or archaea, although PHE degradation has been described among fungi (Haritash et Kaushik 2009; Agrawal, Verma et Shahi 2018). Typically, under aerobic conditions the initial step of the bacterial catabolic pathways starts with the incorporation of molecular oxygen into the aromatic nucleus by a ring hydroxylating dioxygenase (RHD) enzyme system to yield cis dihydrodiol. Cis-dihydrodiol is then rearomatized to a diol intermediate that is further transformed by ring-opening dioxygenases into catechols and other intermediates. Finally, these catechols can be converted to TCA cycle intermediates (Cerniglia 1993; Eaton et Chapman 1992; Mallick, Chakraborty et Dutta 2011). RHD genes are often used as markers of PHE degradation, since they catalyze the first step of the degradation process. They are multicomponent enzyme systems composed of a large alpha subunit and a small beta subunit (Kauppi et al. 1998).The genes coding for the alpha subunit of PAH-RHD form a monophyletic group (Habe et Omori 2003), and were consequently used to design PCR primers for both Gram negative and Gram positive bacteria (Aurélie Cébron et al. 2008), providing a useful way to track bacterial degraders in PAH contaminated environments.

Many microorganisms harbor RHD genes such as members of the genera *Enterobacter* (A. Cébron et al. 2014; Gonzalez et al. 2018; Jha, Annapurna et Saraf 2012) *Sphingomonas* (Li et al. 2019; Ding, Heuer et Smalla 2012; Singha et Pandey 2020), *Burkholderia* (Timm et al. 2016; Aurélie Cébron et al. 2008; Pagé, Yergeau et Greer 2015; Andreolli et al. 2011), *Pseudomonas* (A. Cébron et al. 2014; Wang et al. 2014; Timm et al. 2016; Khan et Bano 2016), *Mycobacterium* (Li et al. 2019; Hormisch et al. 2004; Chen et al. 2016; Guo et al. 2017) and *Polaromonas* (Eriksson, Dalhammar et Mohn 2006; Ding, Heuer et Smalla 2012; Jeon et al. 2004), among others. Moreover, many of these microorganisms have been found to thrive in the rhizosphere of many plants (Thomas, Corre et Cébron 2019; Auffret et al. 2015), including trees, which explains, in part, the success of phytoremediation approaches.

Unravelling the complex microbe-soil-plant relationships within the context of rhizoremediation of PAH contaminated soils remains challenging but was suggested to be key to optimizing phytoremediation to increase its use (Bell et al. 2014; Correa-Garcia et al. 2018). Many recent studies have demonstrated that the diversity of microbial communities has an important role in the degradation prospects in soils (Bell et al. 2016; Bell et al. 2013) and in the rhizosphere of various Salicaceae tree species used for phytoremediation (Yergeau et al. 2015; Bell et al. 2015). For instance, Bell et al. (2016) showed that a soil with a high initial diversity was more efficient at degrading complex hydrocarbon mixture than a soil inoculated with a consortium of confirmed hydrocarbon degraders. Furthermore, under high levels of hydrocarbon contamination, willows grew better when planted in a diverse bulk soil that was not previously planted than in a less diverse rhizospheric soil of a willow that had previously grown well under the same contamination conditions (Yergeau et al. 2015). Taken together these two studies highlight the importance of a diverse initial microbial community over a less diverse pre-selected community for the successful degradation of hydrocarbon in soils. Other studies have shown that the initial community composition also had an importance for the degradation of hydrocarbons in soils. For instance, the initial relative abundance of some of the major families of *Betaproteobacteria* in the soil before contamination were significantly correlated to the amount of diesel degraded across a range of artic soils (Bell et al. 2013). In another study, the total amount of zinc taken up by willows after 16 months of growth was strongly linked to the relative dominance of four fungal taxa in the rhizosphere of the plants after 4 months of growth (Bell 2015). Soil microbiological characteristics appear to have a determining effect on the degradation of hydrocarbons and on the outcome of soil remediation efforts, potentially explaining the variable efficiency of phytoremediation in different soils and, consequently, its low retention as a preferred remediation technique (Mench, 2010).

Here, we hypothesize that differences in the microbiological characteristics of soil (namely, microbial diversity) are a major constrain to PAH rhizoremediation, overriding the rhizosphere effect central to this technology. We aimed at determining the response of the microbial communities of two different soils to contamination with phenanthrene and its subsequent rhizoremediation by poplars. To attain this objective, we designed a pot experiment and sequenced the 16S rRNA gene of bacteria and archaea, the ITS1 region of fungi and the PAH-ring hydroxylating dioxygenase (PAH-RHDα) genes of Gram negative and Gram positive bacteria, and determined the residual concentration of phenanthrene after nine weeks of poplar growth.

## 2. Material and Methods

### 2.1. Experimental design

The experimental design consisted of a full factorial experiment involving three factors: soil contamination, presence of the plant, and soil type. The contamination factor consisted of two levels: soil contaminated with 50 mg kg^-1^ dry soil of phenanthrene (PHE) and uncontaminated soils (CTRL). Plant presence consisted of either a pot without a plant (NP) or a planted pot (P). Finally, the soil type factor consisted of one agricultural and one forest soil. These three factors with two levels each produced 8 treatment combinations. The 12 total replicates per treatment where arranged in four blocks. Pots were randomly distributed inside blocks. A total of 96 pots were placed outside at the Laurentian Forestry Center (Natural Resources Canada, Sainte-Foy, QC, Canada). After planting, the pots were covered with coconut fiber to reduce phenanthrene evaporation and photooxidation. The plants were subsequently connected to an automated irrigation system and received approximately 400 mL of water every day at 8 am.

### 2.2. Soil and biological material

In this experiment, two types of soils were selected: an agricultural soil and a forest soil. The agricultural soil was collected from the 15 cm of the upper soil layer from two sites closely situated within a family farm located in Ste-Famille de l’Île d’Orléans, Quebec, Canada (GPS coordinates site one: 46.941037,-70.974966 and site two: 46.937038, -70.968887). The farm had been alternating crops for more than thirty years with rotations including corn, soy, oat, hay, with fallow every three years. The forest soil used in this experiment was a mix of two different soils. Two thirds (vol/vol) of the forest soil mixture corresponded to soil collected from the upper layer of two closely situated forested plots of the Valcartier forest research station located in Saint-Gabriel-de-Valcartier, Quebec Canada (GPS coordinates for site one: 46.950110, -71.497922 and for site two: 46.951730, -71.497097). Due to the high content of organic matter and low mineral proportion of this forest soil, 1/3 of garden topsoil was incorporated to the forest soil. This was done in order to better compare the two soil type matrices in terms of physical characteristics. The garden topsoil was acquired from a local company (Les developpements Robko Inc., Quebec City, Quebec, Canada). All the soils used were thoroughly homogenized prior to sieving (4 mm mesh) and before setting the experiment. Physicochemical analyses of the experimental potting mixes were performed at AgroEnviroLab, La Pocatière, Quebec, Canada (Suppl Table 1).

Half of each soil was spiked with 50 mg kg^-1^ dry soil of phenanthrene according to the following protocol. One-tenth of the soil was spiked while leaving the rest untreated. Five hundred mg of phenanthrene (Sigma Aldrich) diluted in 100 mL of acetone were applied to 1kg batches of dried 2 mm sieved soil. The soil spiked this way was left outside protected from direct sunlight for 24 h until the acetone was completely evaporated. Then, the spiked batches were thoroughly incorporated into 9 kg of uncontaminated soil in a cement mixer until the potting mix reached homogeneity.

The plants used for the experiments were cuttings harvested from the same adult poplar tree (*Populus balsamifera)*. Cuttings were propagated following the method described in (Rheault et al. 2020). Briefly, cuttings were kept at 4 °C until rooting. Dormancy was disrupted by soaking the cuttings in a rooting mixture for a few weeks, then, planted in 500 g of potting media. Subsequently, between the 19^th^ and the 21^th^ of July 2017, a total of 48 trees were planted into 6 L pots containing approximately 5 kg of either agricultural or forest mix soil, PHE or CTRL. Forty-eight extra pots containing only soil were used for the non-planted treatments (NP).

### 2.3. Sampling and plant trait measurements

The experiment was sampled between the 18^th^ and 20^th^ of September 2017, 9 weeks after planting. Aboveground biomass was clipped, collected and kept at 4 °C until biomass measurements. For the planted pots, the soil adhering to the roots after a vigorous shaking of the root system was considered as “rhizosphere” (P soil). The non-planted soil was collected at a depth of approximately 10 cm (NP soil). Soil samples were kept at 4 °C until transportation to the lab (around 2 h) where they were placed at - 20 °C. Soil samples were used for phenanthrene and microbial community analyses. Harvested plants were weighed fresh and again after 24 h at 60 °C to obtain dry biomass.

### 2.4. Phenanthrene quantification

Soil samples were analyzed for phenanthrene content at the end of the experiment for both NP and P pots. Two phenanthrene measurements were taken per biological replicate. The phenanthrene extraction protocol consisted of the following in-house method developed in the lab.

For sample preparation, approximately 4 g of frozen soil were introduced in a screw cap glass tube. Nine hundred µL of ethyl acetate were added to each sample together with 3 mL of distilled water. Ten ppm of phenanthrene-d_10_ in 100 µL of ethyl acetate was added as the internal standard. Glass tubes containing samples were shaken to thoroughly mix the soil with the organic solvent. Next, the tubes were placed in an ultrasonic bath at 60 kHz for 15 min to detach the remaining phenanthrene from the soil organic matter. Then, the tubes were placed horizontally in a shaker at 300 rpm and left overnight at room temperature. The next day, the tubes were centrifuged at 270 g for 10 min. Afterward, the ethyl acetate phase was recovered and further centrifuged at 5,000 g for 1 min. Lastly, the ethyl acetate containing the phenanthrene was recovered and placed in a GC vial for GC-MS analysis.

Extracts were analyzed for phenanthrene content with a Trace GC Ultra system (Thermo Scientific) equipped with a 30 x 0.25 mm (0.25 µm thickness) DB-5 MS capillary column (Agilent J & W capillary GC) and coupled to a Polaris Q benchtop Ion Trap Mass Spectrometer. The injector and analyzer temperatures were at 250 °C and 350 °C, respectively. The GC-MS temperature program consisted of 2 min hold at 70 °C, increasing temperature to 310 °C at 30 °C min^-1^ followed by 6 min hold at 310 °C. Helium was used as carrier gas at a flow rate of 0.3 mL/min. The injection volume was 3 µL and the MS scan range was from 70-600 m/z.

Calibrations curves for the samples were prepared using phenanthrene as a marker. The phenanthrene was weighed and dissolved in ethyl acetate at concentrations ranging from 1 ppm to 100 ppm. Standard calibration curves were calculated by plotting the peak areas against the corresponding concentration of reference.

### 2.5. DNA isolation and amplicon library preparation

DNA was extracted from up to 250 mg of soil using the DNeasy® Powersoil® DNA Isolation kit (Qiagen) following the protocol provided. The bead-beating step was carried out on a FastPrep®-24 (MP Biomedicals) at 6 m/s for 45 s, twice. Bacterial and archaeal V3-V4 regions of the 16S rRNA gene were amplified using the primer set 515F (5’-GTGCCAGCMGCCGCGGTAA-3’) – 806R (5’-GGACTACHVGGGTWTCTAAT-3’) (Caporaso et al. 2012). The fungal internal transcribed spacer region 1 (ITS1) was amplified using the primers ITS1F (5’-CTTGGTCATTTAGAGGAAGTAA-3’) – 58A2R (5’-CTGCGTTCTTCATCGAT-3’) (Martin and Rygiewicz 2005). The PAH-RHDα genes cluster corresponding to GN bacteria *(nahAc, nahA3, nagAc, ndoB, ndoC2, pahAc, pahA3, phnAc, phnA1, bphAc, bphA1, dntAc and arhA1*) was amplified using the primer set 610F (5’-GAG ATG CATACC ACG TKG GTT GGA-3’) and 916R (5’-AGC TGT TGT TCG GGA AGAYWG TGC MGT T-3’) and the genes cluster corresponding to GP bacteria (*narAa, phdA/pdoA2, nidA/pdoA1, nidA3/fadA1*) was amplified using the primer set 641F (5’-CGG CGC CGA CAA YTT YGT NGG-3’) and 933R (5’-GGG GAA CAC GGTGCC RTG DAT RAA-3’) developed elsewhere (Cébron et al. 2008). All microbial regions were amplified in 25 µL volumes using 19.125 µL of sterile water, 50 nM BSA, 20mM Taq reaction buffer (20mM Tris HCl pH 8.4), 400 nM (0.5 µL at 20 µM) of each primer, 500 nM of MgCl_2_, 200 nM dNTP, 0.25 U Taq DNA polymerase and 1 µL of DNA template (5-10 ng/µL) on a T100 ^TM^ Thermal Cycler (Bio-Rad). Thermal cycling conditions for bacteria and archaea were as follows: initial denaturation at 95 °C for 5 min; 35 cycles at 95 °C for 30 s, 55 °C for 30 s, 72 °C for 1 min, and a final elongation phase at 72 °C for 5 minutes. For fungi, a similar protocol was used except that the annealing temperature was at 59 °C and 30 cycles were performed. PAH-RHDα genes also had different annealing temperatures set at 57 °C and 54 °C for GN and GP primer sets, respectively for a total of 30 cycles. Following amplification, PCR products were cleaned using 16 µL of magnetic beads (Agencourt AMPure XP, Beckman Coulter Life Science) following the Illumina’s protocol 16S Metagenomic Sequencing Library preparation guide (Part #15044223 Rev. B). Then, 400 nM of each Nextera XT unique index primers (2 µL at 5 µM) were added to 5 µL of our PCR product, in a PCR mix containing 50 nM BSA, 20 mM Taq reaction buffer (20mM Tris HCl pH 8.4), 500 nM of MgCl_2_, 200 nM dNTP, 0.25 U Taq DNA polymerase and 12.175 µL of sterile water. The PCR product was tagged under the following thermal cycling conditions: 95 °C initial denaturation phase for 5 min, followed by 8 cycles of denaturation at 95 °C for 30 s, annealing at 55 °C for 30 s, elongation at 68 °C for 30 s, and a final elongation phase at 68 °C during 5 min. The indexed amplicons were purified using magnetic beads as described above, and quantified the DNA using the Quant-iT^TM^ Picogreen dsDNA Assay kit (Invitrogen). The libraries were combined in an equimolar ratio and sent them for 2 x 250 bp pair-end sequencing on an Illumina Miseq at The Genome Quebec Service Center (Montréal, QC, Canada).

### 2.6. Quantification of PAH degrading bacteria (qPCR)

The real-time quantitative PCR (qPCR) was conducted on a Stratagene Mx3005P qPCR system (Agilent Technologies), associated with the corresponding software MxPro Mx3005P (v4.10; Agilent). The qPCR reactions were performed using the primers designed by Cébron et al. (2008) in 20 µL total volume containing 1x iTaq universal SYBR® Green reaction mix supplemented with 300 µM of each primer (Integrated DNA Technologies™) and 5 µL of DNA template at a concentration around 5-10 ng µL^-1^ or distilled sterile water (negative no template control).

The amplifications were carried out following the protocol provided in Cébron et al. (2008) with some modifications. Briefly, the first step consisted of denaturation at 95 °C for 5 min followed by 40 cycles of denaturation at 95 °C for 30 s, annealing for 35 s at either 57 °C (GN) or 54 °C (GP) and an elongation step at 72 °C for 75 s, after which the SYBR Green signal intensities were measured. At the end of the run, a melting curve analysis was performed where signal intensities were measured at 0.5 °C temperature increment every 5 s from 51 to 95 °C. Standards for each gene were made from 10-fold dilutions of linearized plasmid containing the gene fragment of interest, cloned from amplified soil DNA (Yergeau et al. 2009).

At the end of the qPCR run, the threshold line was automatically defined within the logarithmic increase phase of the acquired fluorescence data. The Ct values were assessed for all the samples and the gene copy numbers were deducted from the standard curve based on the Ct value. Efficacy of the qPCRs were of 89.3% (standard curve R^2^ = 0.998) and 92% (R^2^ = 0.995) for PAH-RHDα GN genes and 90.6% (R^2^ = 0.995) and 85.8% (R^2^ = 0.999) for PAH-RHDα GP genes.

### 2.7. Bioinformatic analysis

Sequences were analyzed AmpliconTagger (Tremblay et Yergeau, 2019). Briefly, raw reads were scanned for sequencing adapters and PhiX spike-in sequences. Remaining reads were filtered based on quality (Phred) score and remaining sequences were dereplicated/clustered at 97% identity (16S rRNA and ITS genes) and then processed for generating Operational Taxonomic Units (OTUs) (DADA2 v1.12.1) (PMID:27214047). Chimeras were removed with DADA2’s internal removeBimeraDeNovo(method=”consensus”) method followed by UCHIME reference (Edgar et al., 2011). OTUs for which abundance across all samples were lower than 5 were discarded. A global read count summary throughout the pipeline steps is provided in Suppl. Table 5 for all datasets. OTUs were assigned a taxonomic lineage with the RDP classifier (PMID: 17586664) using an in-house training set containing the complete Silva release 138 database (PMID:23193283) supplemented with eukaryotic sequences from the Silva database and a customized set of mitochondria, plasmid and bacterial 16S sequences. For ITS OTUs, a training set containing the Unite DB was used (sh_general_release_s_04.02.2020 version). The RDP classifier assigns a score (0 to 1) to each taxonomic depth of each OTU. Each taxonomic depth having a score ≥ 0.5 were kept to reconstruct the final lineage. Taxonomic lineages were combined with the cluster abundance matrix obtained above to generate a raw OTU table, from which a bacterial organisms OTU table was generated.

Custom RDP classifier training sets were generated for both PAH-RHD GP and GN amplicon data types. Raw reads were processed as described above up to the quality filtering step. Then, remaining sequences were dereplicated/clustered at 100%, thus processes for generating Amplicon Sequence Variants (ASVs) (DADA2 v1.12.1) (PMID:27214047).

Each ASV were blasted (BLASTn) against the NCBI nt database (downloaded as of 21^st^ February 2020) with –max_target_seqs set to 20. Blast output was filtered to keep hits that had an e-value <= 1e-20, alignment length of at least 100 bp and alignment percentage of at least 60 percent. Taxonomic lineages of each filtered blast hit were fetched from the NCBI taxonomy database (version downloaded on December 4th 2019). RDP training sets were generated as described (https://github.com/jtremblay/RDP-training-sets).

### 2.8. Statistical analysis

Statistical analyses were performed in R (v.3.5.0)(R Core Team 2020), with basic analysis performed with the stats package (R Core Team 2020). Normality and homoscedasticity of data were calculated with Shapiro-Wilk and Bartlett tests, respectively. When distributions did not attain normality, variables were log transformed or converted using a Box-Cox transformation using the MASS package (Venables et Ripley 2002). The differences in plant trait results and phenanthrene concentration were assessed through univariate analysis of variance using ANOVA followed by *post hoc* Tukey HSD tests. When transformations did not yield linearity, Kruskal-Wallis analysis of the variance was performed instead, with the Dunn tests performed for multiple comparisons. Univariate analysis of relative abundance of genera were performed with ANOVA test with White’s correction for heteroscedastic data using the car package (Fox et Weisberg 2019). Differences were considered significant at the 0.05 level.

Shannon diversity index and Species Richness were calculated with the otuSummary package (Yang 2020).

Principal coordinate analyses (PCoA) were performed to assess differences in community composition and between treatments using normalized OTU or ASV tables with the Bray-Curtis dissimilarity index calculated with the vegan package (Oksanen et al. 2019). PERMANOVA analyses tested the interaction between treatments in the microbial communities through 999 permutations using the Adonis function from vegan. All graphs were created with the ggplot2 package (Wickham et al. 2019). Finally, correlation analysis where performed with the stats package.

## 3. Results

### 3.1. Plant morphological response and phenanthrene degradation

All the 48 poplar cuttings survived throughout the experiment and were included in the analysis. Poplar growth was evaluated as the plant aboveground biomass but neither soil type nor contamination had a significant impact on this variable (Fig. 1a).

**Figure 1.**
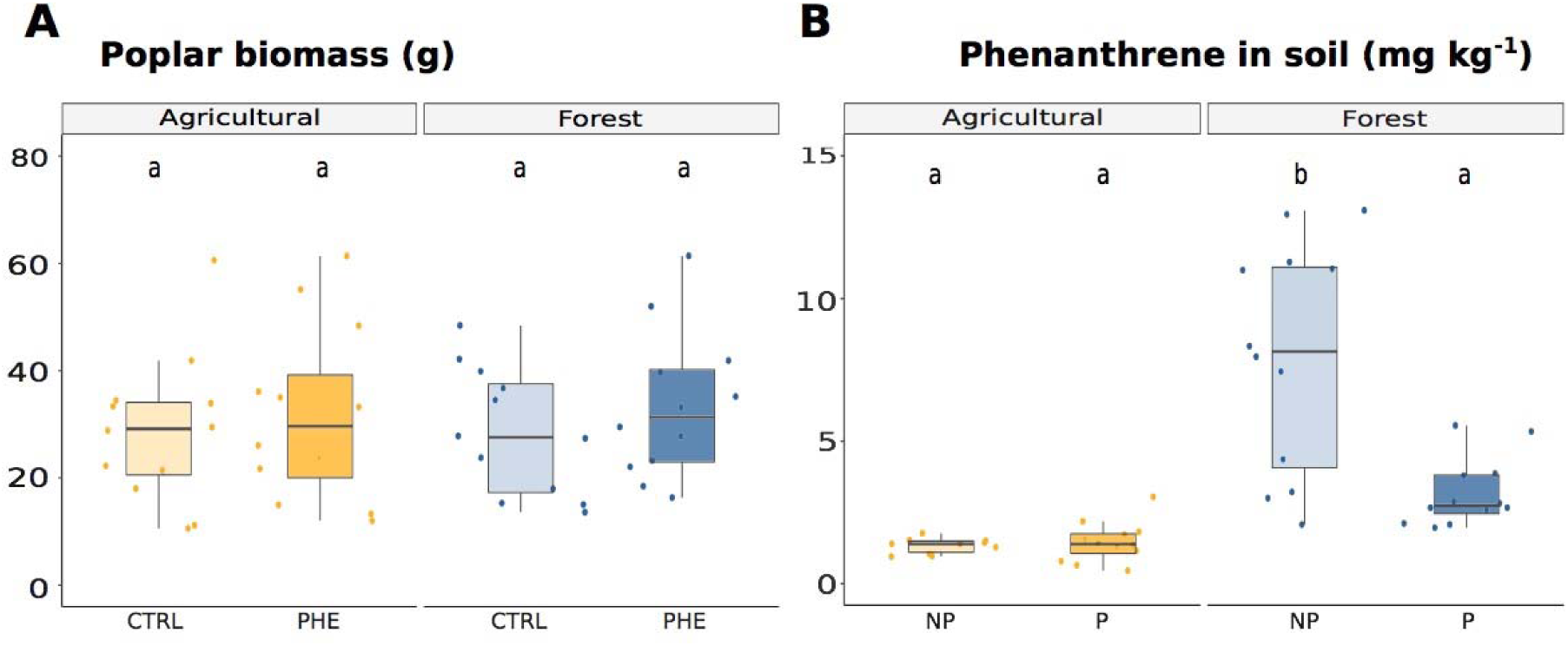
Boxplots of poplar biomass and phenanthrene presence. A) Dried biomass of poplars grown in agricultural and forest soil contaminated with phenanthrene (PHE) or not (CTRL); **B**) Quantification of phenanthrene in agricultural and forest soils after a nine weeks of poplar growth in pots (P, for planted) compared to non-planted pots (NP). Different letters denote significant differences found with the Tukey HSD post-hoc test (n = 12).

Phenanthrene degradation was measured as the quantity of remaining phenanthrene after nine weeks of incubation and was expressed as absolute mg kg^-1^ retrieved in soil. The soil type (F = 79.508, p <0.001), plant presence (F = 9.442, p = 0.004) and the interaction term (F = 8.166, p = 0.007) were all significant (Fig. 1b). The interaction term revealed that the effect of soil type depended on plant presence. More precisely, in agricultural soils, both P and NP pots showed similar degradation, with 1.33 and 1.45 mg kg^-1^ of phenanthrene left in NP and P pots, respectively. In contrast, after nine weeks the non-planted forest soils contained significantly more phenanthrene than the planted forest soils (7.97 vs. 3.19 mg kg^-1^ of phenanthrene; Fig. 1b).

### 3.2. Microbial community response

#### 3.2.1. Fungi

##### Fungal diversity

Shannon diversity index for fungi ranged from 1.599 to 3.646 (Fig. 2a). The Kruskal-Wallis test showed significant differences among treatments (χ² = 17.779, p-value = 0.013). The Dunn test showed that this difference was due to significantly higher Shannon diversity in the P CTRL forest soils as compared to the NP CTRL and NP PHE agricultural soils and to the NP PHE forest soil (Fig. 2a – Shannon index). The same trends were visible for species richness, but this was not statistically significant (Fig. 2a – Species richness).

**Figure 2.**
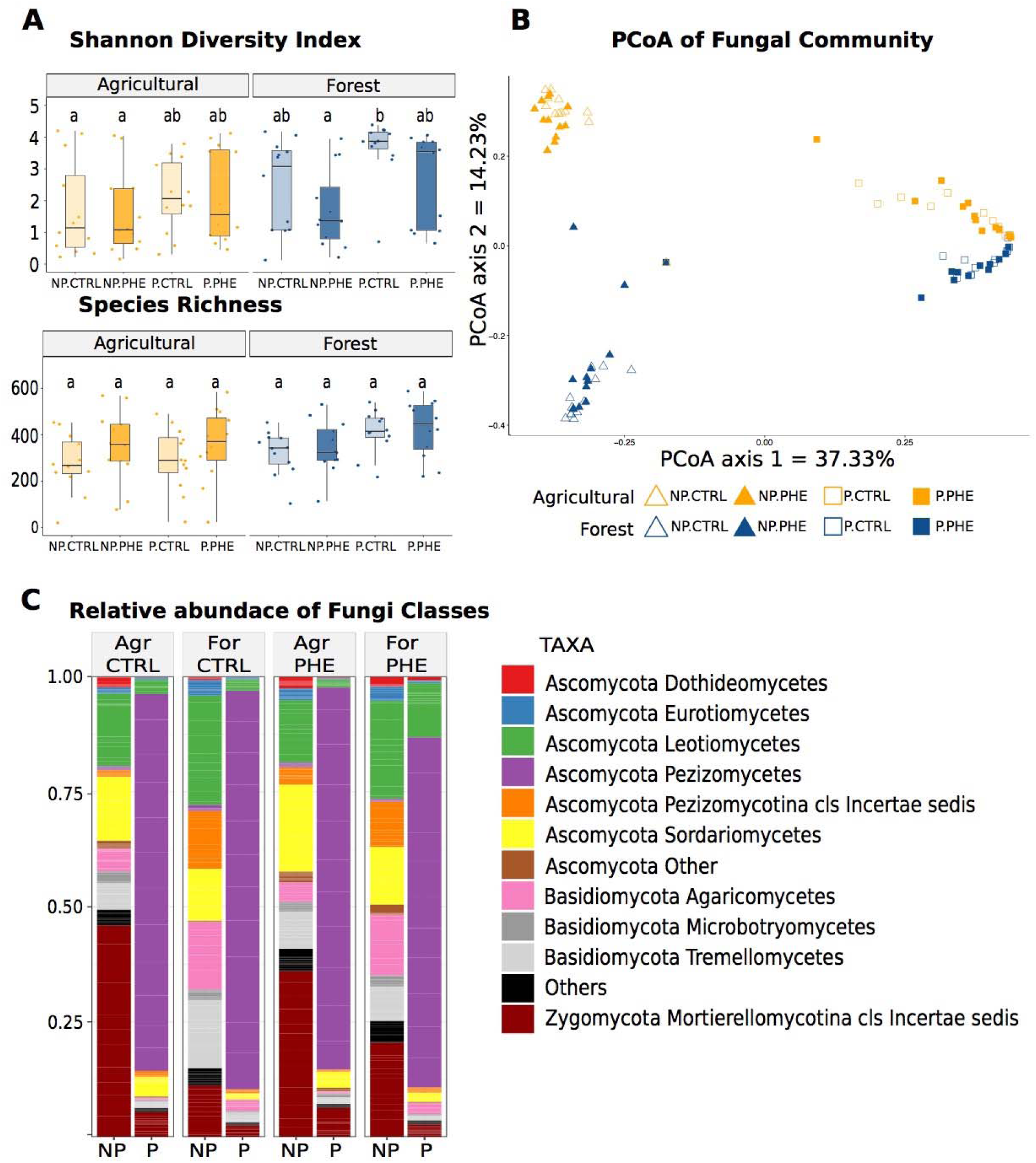
Summary of the fungal community diversity, structure and composition based on OTUs of the ITS region. **A**) Boxplots of Shannon diversity index and Species richness (as observed number of OTUs) by treatment. Different letters denote significant differences found with the Tukey HSD *post-hoc* test. **B**) Principal coordinates analysis (PCoA) based on Bray-Curtis dissimilarity of the relative abundance of fungal OTUs showing the effects of contamination, soil type and plant presence on the fungal community structure. **C**) Fungal community composition at the class level. Only taxa with a relative abundance above 0.01 are shown. Values are averaged across treatments (n = 12). Legend: NP: non planted pots. P: planted pots. CTRL: non contaminated pots. PHE: pots contaminated with 50 mg kg ^-1^ of phenanthrene.

##### Fungal community structure

Analysis of the beta diversity with the PCoA (Fig. 2b) based on relative abundance showed that the fungal community structured differed between plant presence and soil types. The main factor was plant presence (R^2^ = 49.7%, p = 0.001), followed by the soil type (R^2^ = 14.5%, p = 0.001). However, the main plant effect was modulated by the interaction with the soil type factor (R^2^ = 5.0%, p = 0.001). However, contamination, either as a main effect or its interaction with other factors had no significant influence on the fungal community structure (Table 1).

**Table 1.** Summary of permutational multivariate analysis of variance (PERMANOVA) examining differences in microbial composition based on the OTUs of the ITS region for fungi and 16S RNA gene for total bacteria and ASVs for GN and GP PAH-RHDα genes for bacterial degraders. Values in bold indicate significant effects. Df, degrees of freedom. SumsOfSqs, sum of squares. MeanSqs, Mean squares. Pr(>F), p-value.

| **Fungi** |  |  |  |  |  |  |
| --- | --- | --- | --- | --- | --- | --- |
| **PERMANOVA** | *Df* | *SumsOfSqs* | *MeanSqs* | *F Model* | *R<sup>2</sup>* | *Pr(>F)* |
| **soil** | **1** | **3.938** | **3.938** | **44.745** | **0.145** | **0.001** |
| **contamination** | 1 | 0.109 | 0.109 | 1.234 | 0.004 | 0.26 |
| **plant** | **1** | **13.526** | **13.526** | **153.690** | **0.497** | **0.001** |
| **soil:contamination** | 1 | 0.174 | 0.174 | 1.977 | 0.006 | 0.101 |
| **soil:plant** | **1** | **1.353** | **1.353** | **15.377** | **0.050** | **0.001** |
| **contamination:plant** | 1 | 0.148 | 0.148 | 1.678 | 0.005 | 0.157 |
| **soil:contamination:plant** | 1 | 0.197 | 0.197 | 2.241 | 0.007 | 0.078 |
| **Residuals** | 88 | 7.745 | 0.088 | NA | 0.285 | NA |
| **Total** | 95 | 27.189 | NA | NA | 1.000 | NA |
| **Bacteria** |  |  |  |  |  |  |
| **soil** | **1** | **12.348** | **12.348** | **115.521** | **0.512** | **0.001** |
| **contamination** | **1** | **0.530** | **0.530** | **4.955** | **0.022** | **0.004** |
| **plant** | **1** | **0.572** | **0.572** | **5.353** | **0.024** | **0.003** |
| **soil:contamination** | **1** | **0.451** | **0.451** | **4.221** | **0.019** | **0.009** |
| **soil:plant** | **1** | **0.460** | **0.460** | **4.301** | **0.019** | **0.005** |
| **contamination:plant** | 1 | 0.165 | 0.165 | 1.542 | 0.007 | 0.150 |
| **soil:contamination:plant** | 1 | 0.169 | 0.169 | 1.583 | 0.007 | 0.145 |
| **Residuals** | 88 | 9.406 | 0.107 | NA | 0.390 | NA |
| **Total** | 95 | 24.101 | NA | NA | 1.000 | NA |
| **PAH-RHDα GN genes** |  |  |  |  |  |  |
| **soil** | **1** | **4.341** | **4.341** | **14.912** | **0.119** | **0.001** |
| **contamination** | **1** | **2.622** | **2.622** | **9.007** | **0.072** | **0.001** |
| **plant** | 1 | 0.283 | 0.283 | 0.971 | 0.008 | 0.457 |
| **soil:contamination** | **1** | **3.091** | **3.091** | **10.618** | **0.085** | **0.001** |
| **soil:plant** | 1 | 0.193 | 0.193 | 0.663 | 0.005 | 0.811 |
| **contamination:plant** | 1 | 0.228 | 0.228 | 0.782 | 0.006 | 0.659 |
| **soil:contamination:plant** | 1 | 0.293 | 0.293 | 1.005 | 0.008 | 0.401 |
| **Residuals** | 86 | 25.034 | 0.291 | NA | 0.688 | NA |
| **Total** | 94 | 36.402 | NA | NA | 1.000 | NA |
| **PAH-RHDα GP genes** |  |  |  |  |  |  |
| **soil** | **1** | **4.900** | **4.900** | **16.970** | **0.146** | **0.001** |
| **contamination** | **1** | **1.230** | **1.230** | **4.261** | **0.037** | **0.001** |
| **plant** | **1** | **0.644** | **0.644** | **2.232** | **0.019** | **0.014** |
| **soil:contamination** | 1 | 0.340 | 0.340 | 1.176 | 0.010 | 0.176 |
| **soil:plant** | 1 | 0.417 | 0.417 | 1.446 | 0.012 | 0.082 |
| **contamination:plant** | 1 | 0.441 | 0.441 | 1.528 | 0.013 | 0.08 |
| **soil:contamination:plant** | 1 | **0.670** | **0.670** | **2.322** | **0.020** | **0.01** |
| **Residuals** | 85 | 24.54 | 24.54 | NA | 0.733 | NA |
| **Total** | 93 | 36.402 | NA | NA | 1.000 | NA |

More specifically, while the fungal communities found in the NP pots diverged significantly from one another depending on the soil type, these communities tended to converge in the P pots. The PERMANOVA analysis confirmed this visual interpretation, with plant presence representing the main source of variation, followed by first order effect of soil type and the interaction of both factors (Table 1).

##### Fungal community composition

A total of 3,099 fungal OTUs were retained after the bioinformatic quality filtering pipeline. As for the community structure, the fungal community composition was principally influenced by plant presence. As such, the rhizosphere (P) of both PHE and CTRL in forest and agricultural soils were vastly dominated by fungi from the genus *Sphaerosporella* (Class *Pezizomycetes*; Fig. 2c and Suppl. Table 2). The NP pots of both types of soil showed mixed community composition, strongly distinct from the P pots. The lower diversity of fungi in the agricultural soil could be explained by the prevalence of *Mortierellomycotina* (Fig. 2c and Suppl. Table 2), that accounted for around 45% and 35% of total OTUs in CTRL and PHE agricultural NP pots, respectively. In contrast, the presence of *Mortierellomycotina* in forest soils was lower than 20%, whereas *Basidiomycota* (*Tremellomycetes* and *Agaricomycetes* classes) were significantly more abundant (Fig 2c and Suppl. Table 2). Within the *Ascomycota* phylum, *Leotiomycetes* were also significantly more abundant in forest than in agricultural soils (Suppl. Table 2).

Some significant but weak correlations were found among some fungal OTUs and the plant biomass. The top 10 OTUs had a Pearsons’s correlation coefficient r ranging from 0.395 to 0.331, and the OTUs belonged mainly to the *Ascomycota* phylum.

The significant correlations found between fungi and phenanthrene concentration in soils at the end of the experiment were stronger (Pearsons’s correlation coefficient r between 0.834-0.744). The top 10 OTUs presenting positive correlations belonged to the genera: *Aspergillus*, *Acrodontium, Imleria, Postia*, *Umbelopsis*, *Acrophialophora*, *Wilcoxina,* an unclassified *Basidiomycota* and *Leptodontidium* (Suppl. Table 3).

#### 3.2.2. Bacteria

##### Bacterial diversity

A total of 45 archaeal OTUs were found in the soil samples representing a combined relative abundance of less than 0.1%, therefore they were not retained for the analyses.

In terms of bacterial diversity, there were no significant differences between treatments although slightly higher values were found in NP PHE soils for the Shannon index (Fig. 3a - Shannon index). For the species richness, the forest soils seemed to present higher number of OTUs in PHE pots while the agricultural soils showed higher OTU numbers in CTRL soils (Fig. 3a - Species richness), but, again, this difference was not significant.

**Figure 3.**
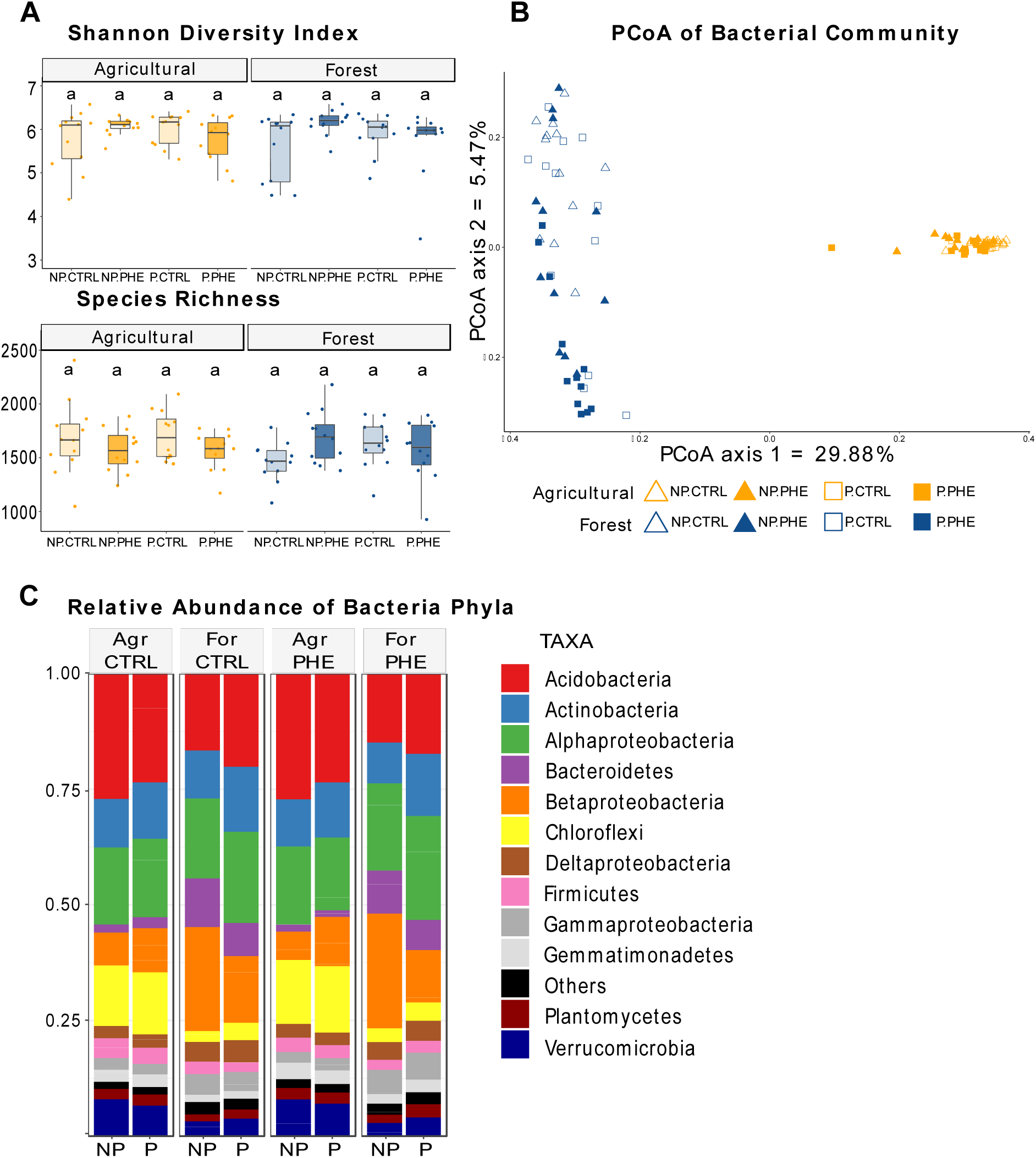
Summary of the bacterial community diversity, structure and composition based on OTUs of the 16 rRNA gene. **A**) Boxplots of Shannon diversity index and Species richness (as observed number of OTUs) by treatment. Different letters denote significant differences found with the Tukey HSD *post-hoc* test. **B**) Principal coordinates analysis (PCoA) based on Bray-Curtis dissimilarity of the relative abundance of bacterial OTUs showing the effects of contamination, soil type and plant presence on the bacterial community structure. **C**) Bacterial community composition at the class level. Only taxa with a relative abundance above 0.01 are shown. Values are averaged across treatments (n = 12). Legend: NP: non planted pots. P: planted pots. CTRL: non contaminated pots. PHE: pots contaminated with 50 mg kg ^-1^ of phenanthrene.

##### Bacterial community structure

The PCoA based on relative abundance (Fig. 3b) along with the PERMANOVA analysis (Table 1), showed that the soil type was the main factor shaping the bacterial community (R^2^ = 51.2%, p = 0.001), separating agricultural from forest soil samples along the first axis, in sharp contrast to what we observed in the fungal community structure. Moreover, plant presence (R^2^ = 2.4%, p = 0.003) and the contamination status (R^2^ = 2.2%, p = 0.004), had minor but significant main effects and also modulated the effect of soil type through significant interaction terms (R^2^ = 1.9%, p = 0.005 and p = 0.009, respectively; Table 1). These interaction effects are visible in Fig. 3b, as the bacterial communities found in the agricultural soils all clustered together, whereas the bacterial communities of the forest soil were also clustering loosely based on soil contamination and plant presence on the second axis of the ordination.

##### Bacterial community composition

A total of 5,716 OTUs were retained from the 16S rRNA gene sequences. From those, the vast majority belonged to Bacteria, and only 45 OTUs were Archaea.

The 5,671 bacterial OTUs were classified in 35 phyla, although the majority of OTUs belonged to *Proteobacteria*, *Actinobacteria*, *Acidobacteria*, *Bacteroidetes*, *Chloroflexi*, *Firmicutes*, *Gemmatimonadetes Planctomycetes* and *Verrucomicrobia*. Even though agricultural and forest soils presented very different community structures (Fig. 3b), the composition at the phylum level was relatively homogeneous among treatments (Fig 3c), with a few significant differences in terms of the overall relative abundance of phyla (Suppl. Table 4).

In contrast, major differences were observed in the relative abundance at genus levels (Table 2). Compared to forest soils, agricultural soils had significantly higher relative abundance of the following genera: unclassified uncultured *Acidobacteriaceae* Subgroup 1 (*Acidobacteria*, mean of 14.84% vs 2.52%), HSB OG53-F07FA (*Chloroflexi*, 8.07% vs 0.12%), DA101 soil group FA (*Verrucomicrobia*, 5.06% vs 0.36%), Candidatus *Solibacter* (*Acidobacteria*, 2.40% vs 0.94%) *Sphingomonas* (*Alphaproteobacteria*, 3.51% vs 2.50%) and *Gemmatimonas* (*Gemmatomonadetes* 2.40% vs 1.27%), the latest being mostly absent in the forest soils. In contrast, the forest soils had significantly higher relative abundances of the genera: *Ramlibacter* (*Betaproteobacteria* 1.11% vs 4.66%), *Geothrix* (*Acidobacteria*, 0.02% vs 5.61%) and *Variibacter* (*Alphaproteobacteria*, 2.88% vs 4.6%).

**Table 2.** Summary of the three-way analysis of the variance (ANOVA) – with White’s correction for heteroscedasticity on the relative abundance of the 16S rRNA gene OTUs identified at the genus level of bacteria. Values in bold indicate significant effects. Df, degrees of freedom. SumsOfSqs, sum of squares. MeanSqs, Mean squares. Pr(>F), p-value.

| ***Acidobacteria* Subgroup 2CL** |  |  |  |
| --- | --- | --- | --- |
|  | *Df* | *F* | *Pr(>F)* |
| **soil** | **1.000** | **6.402** | **0.013** |
| contamination | 1.000 | 1.200 | 0.276 |
| plant | 1.000 | 0.333 | 0.565 |
| soil:contamination | 1.000 | 0.232 | 0.632 |
| **soil:plant** | **1.000** | **18.052** | **<0.001** |
| contamination:plant | 1.000 | 2.681 | 0.105 |
| soil:contamination:plant | 1.000 | 0.236 | 0.629 |
| Residuals | 88.000 |  |  |
| ***Actinobacteria Acidothermus*** |  |  |  |
|  | *Df* | *F* | *Pr(>F)* |
| **soil** | **1.000** | **202.047** | **<0.001** |
| contamination | 1.000 | 2.195 | 0.142 |
| **plant** | **1.000** | **54.144** | **<0.001** |
| soil:contamination | 1.000 | 0.606 | 0.438 |
| **soil:plant** | **1.000** | **8.664** | **0.004** |
| contamination:plant | 1.000 | 0.034 | 0.855 |
| soil:contamination:plant | 1.000 | 2.662 | 0.106 |
| Residuals | 88.000 |  |  |
| ***Gammaproteobacteria Aquincola*** |  |  |  |
|  | *Df* | *F* | *Pr(>F)* |
| soil | 1.000 | 1.083 | 0.301 |
| **contamination** | **1.000** | **29.161** | **<0.001** |
| plant | 1.000 | 2.822 | 0.097 |
| soil:contamination | 1.000 | 2.972 | 0.088 |
| **soil:plant** | **1.000** | **18.062** | **<0.001** |
| contamination:plant | 1.000 | 0.634 | 0.428 |
| soil:contamination:plant | 1.000 | 0.034 | 0.853 |
| Residuals | 88.000 |  |  |
| ***Actinobacteria Pseudarthrobacter*** |  |  |  |
|  | *Df* | *F* | *Pr(>F)* |
| **soil** | **1.000** | **23.154** | **<0.001** |
| contamination | 1.000 | 0.165 | 0.685 |
| **plant** | **1.000** | **6.909** | **0.010** |
| **soil:contamination** | **1.000** | **6.797** | **0.011** |
| soil:plant | 1.000 | 3.357 | 0.070 |
| contamination:plant | 1.000 | 2.648 | 0.107 |
| soil:contamination:plant | 1.000 | 0.531 | 0.468 |
| Residuals | 88.000 |  |  |

| ***Acidobacteria Geothrix*** |  |  |  |
| --- | --- | --- | --- |
|  | *Df* | *F* | *Pr(>F)* |
| **soil** | **1.000** | **46.073** | **<0.001** |
| contamination | 1.000 | 1.607 | 0.208 |
| plant | 1.000 | 1.799 | 0.183 |
| **soil:contamination** | **1.000** | **4.526** | **0.036** |
| soil:plant | 1.000 | 1.005 | 0.319 |
| contamination:plant | 1.000 | 0.004 | 0.947 |
| soil:contamination:plant | 1.000 | 0.710 | 0.402 |
| Residuals | 88.000 |  |  |
| ***Betaproteobacteria Massilia*** |  |  |  |
|  | *Df* | *F* | *Pr(>F)* |
| soil | 1.000 | 0.038 | 0.847 |
| **contamination** | **1.000** | **6.642** | **0.012** |
| plant | 1.000 | 1.497 | 0.224 |
| soil:contamination | 1.000 | 1.138 | 0.289 |
| soil:plant | 1.000 | 3.685 | 0.058 |
| contamination:plant | 1.000 | 0.491 | 0.486 |
| soil:contamination:plant | 1.000 | 2.378 | 0.127 |
| Residuals | 88.000 |  |  |
| ***Betaproteobacteria Polaromonas*** |  |  |  |
|  | *Df* | *F* | *Pr(>F)* |
| **soil** | **1.000** | **71.821** | **<0.001** |
| contamination | 1.000 | 0.612 | 0.436 |
| plant | 1.000 | 0.173 | 0.679 |
| **soil:contamination** | **1.000** | **4.101** | **0.046** |
| **soil:plant** | **1.000** | **10.727** | **0.002** |
| contamination:plant | 1.000 | 1.798 | 0.183 |
| **soil:contamination:plant** | **1.000** | **10.349** | **0.002** |
| Residuals | 88.000 |  |  |
| ***Alphaproteobacteria Variibacter*** |  |  |  |
|  | *Df* | *F* | *Pr(>F)* |
| **soil** | **1.000** | **67.316** | **<0.001** |
| contamination | 1.000 | 0.057 | 0.811 |
| **plant** | **1.000** | **9.031** | **0.003** |
| soil:contamination | 1.000 | 1.088 | 0.300 |
| **soil:plant** | **1.000** | **8.056** | **0.006** |
| contamination:plant | 1.000 | 0.200 | 0.656 |
| soil:contamination:plant | 1.000 | 2.349 | 0.129 |
| Residuals | 88.000 |  |  |
| ***Alphaproteobacteria Sphingomonas*** |  |  |  |
|  | *Df* | *F* | *Pr(>F)* |
| **soil** | **1.000** | **8.023** | **0.006** |
| **contamination** | **1.000** | **5.332** | **0.023** |
| **plant** | **1.000** | **12.001** | **<0.001** |
| **soil:contamination** | **1.000** | **5.730** | **0.019** |
| soil:plant | 1.000 | 1.643 | 0.203 |
| contamination:plant | 1.000 | 0.407 | 0.525 |
| soil:contamination:plant | 1.000 | 1.358 | 0.247 |
| Residuals | 88.000 |  |  |
| ***Betaproteobacteria Ramlibacter*** |  |  |  |
|  | *Df* | *F* | *Pr(>F)* |
| **soil** | **1.000** | **133.801** | **<0.001** |
| **contamination** | **1.000** | **29.482** | **<0.001** |
| plant | 1.000 | 0.486 | 0.488 |
| **soil:contamination** | **1.000** | **4.124** | **0.045** |
| **soil:plant** | **1.000** | **4.620** | **0.034** |
| contamination:plant | 1.000 | 3.097 | 0.082 |
| soil:contamination:plant | 1.000 | 0.673 | 0.414 |
| Residuals | 88.000 |  |  |
| ***Gemmatimonadetes Gemmatimonas*** |  |  |  |
|  | *Df* | *F* | *Pr(>F)* |
| **soil** | **1.000** | **57.249** | **<0.001** |
| **contamination** | **1.000** | **9.794** | **0.002** |
| plant | 1.000 | 1.408 | 0.239 |
| soil:contamination | 1.000 | 0.023 | 0.879 |
| soil:plant | 1.000 | 1.705 | 0.195 |
| contamination:plant | 1.000 | 0.266 | 0.608 |
| soil:contamination:plant | 1.000 | 2.930 | 0.090 |
| Residuals | 88.000 |  |  |
| ***Acidobacteria Candidatus Solibacter*** |  |  |  |
|  | *Df* | *F* | *Pr(>F)* |
| **soil** | **1.000** | **299.754** | **<0.001** |
| contamination | 1.000 | 1.219 | 0.273 |
| **plant** | **1.000** | **28.450** | **<0.001** |
| soil:contamination | 1.000 | 0.019 | 0.892 |
| **soil:plant** | **1.000** | **10.756** | **0.001** |
| contamination:plant | 1.000 | 0.290 | 0.591 |
| soil:contamination:plant | 1.000 | 0.392 | 0.533 |
| Residuals | 88.000 |  |  |
| ***Verrucomicrobia DA101 soil groupFA*** |  |  |  |
|  | *Df* | *F* | *Pr(>F)* |
| **soil** | **1.000** | **603.476** | **<0.001** |
| contamination | 1.000 | 3.051 | 0.084 |
| plant | 1.000 | 0.018 | 0.894 |
| soil:contamination | 1.000 | 0.228 | 0.634 |
| **soil:plant** | **1.000** | **19.820** | **<0.001** |
| contamination:plant | 1.000 | 0.037 | 0.849 |
| soil:contamination:plant | 1.000 | 0.643 | 0.425 |
| Residuals | 88.000 |  |  |
| ***Chloroflexi HSB OF53 F07FA*** |  |  |  |
|  | *Df* | *F* | *Pr(>F)* |
| **soil** | **1.000** | **580.036** | **<0.001** |
| contamination | 1.000 | 0.273 | 0.603 |
| **plant** | **1.000** | **20.951** | **<0.001** |
| soil:contamination | 1.000 | 0.170 | 0.681 |
| soil:plant | 1.000 | 1.232 | 0.270 |
| contamination:plant | 1.000 | 2.152 | 0.146 |
| soil:contamination:plant | 1.000 | 0.000 | 0.993 |
| Residuals | 88.000 |  |  |
| ***Acidobacteria* uncultured *Acidobacteriaceae* Subgroup 1** |  |  |  |
|  | *Df* | *F* | *Pr(>F)* |
| **soil** | **1.000** | **605.363** | **<0.001** |
| contamination | 1.000 | 0.201 | 0.655 |
| **plant** | **1.000** | **11.038** | **0.001** |
| soil:contamination | 1.000 | 1.004 | 0.319 |
| **soil:plant** | **1.000** | **11.469** | **0.001** |
| contamination:plant | 1.000 | 3.830 | 0.054 |
| soil:contamination:plant | 1.000 | 0.019 | 0.891 |
| Residuals | 88.000 |  |  |

Also, the forest soils presented some particularities linked to the effects of the contamination and plant presence that were absent in agricultural soils, and that explained the different community structure captured in the PCoA (Fig. 3b). For instance, regarding the effect of plant presence, *Polaromonas* (*Betaproteobacteria*), *Aquincola* (*Betaproteobacteria*) and *Acidothermus* (*Actinobacteria*) were nearly absent from the P pots, while the *Acidobacteria* subgroup 2CL was significantly more abundant in the planted forest soils as compared to the NP soils (Table 2). Moreover, the significant effect of the interaction between plant presence and contamination, was greatly explained by the higher mean relative abundance of bacteria from the genus *Polaromonas* (*Betaproteobacteria*) in NP PHE forest soils (10.95%) compared to the P PHE of forest soils (1.46%), while being nearly absent in CTRL pots (Table 2).

As it was for fungi, the correlations between bacterial OTUs’ relative abundances and the biomass of the plant were weak. The top ten significant positive correlations had a Pearsons’s correlation coefficient r ranging from 0.476 to 0.414. These OTUs were diversely classified as *Alphaproteobacteria*, *Chloroflexi*, *Acidobacteria*, *Armatimonadetes*, *Actinobacteria*, and *Gammaproteobacteria*. The strongest positive correlation involved an OTU from the genus *Sphingomonas*.

The OTU with the strongest positive correlation with the amount of phenanthrene in soil was a member of the *Polaromonas* genus (Pearsons’s correlation coefficient r = 0.923) which was one of the more abundant genera found in NP PHE forest soils. Moreover, the top ten highly correlated OTUs all presented very high correlation coefficients (Pearsons’s correlation coefficient r ranging from 0.880 to 0.824) and were mostly present in the NP pots (Suppl. Table 3).

Alternatively, the strongest negative correlations between OTUs and phenanthrene concentrations in soil, were mostly for OTUs well represented in the agricultural soil (Suppl. Table 3).

#### 3.2.3. Gram-Negative bacteria degraders

##### PAH-RHDα GN gene diversity

Three phyla were found in the PAH-RHDα GN gene dataset: *Bacteroidetes*, *Firmicutes* and *Proteobacteria*. When looking at the Shannon diversity index, the mean values ranged from 2.442 to 3.801 with significant differences among treatments (Fig. 4a). For P and NP agricultural soils, PAH-RHDα GN Shannon diversity was significantly higher in the PHE soils compared to their corresponding CTRL (Fig. 4a). This pattern was similar for forest soils, but not statistically significant.

**Figure 4.**
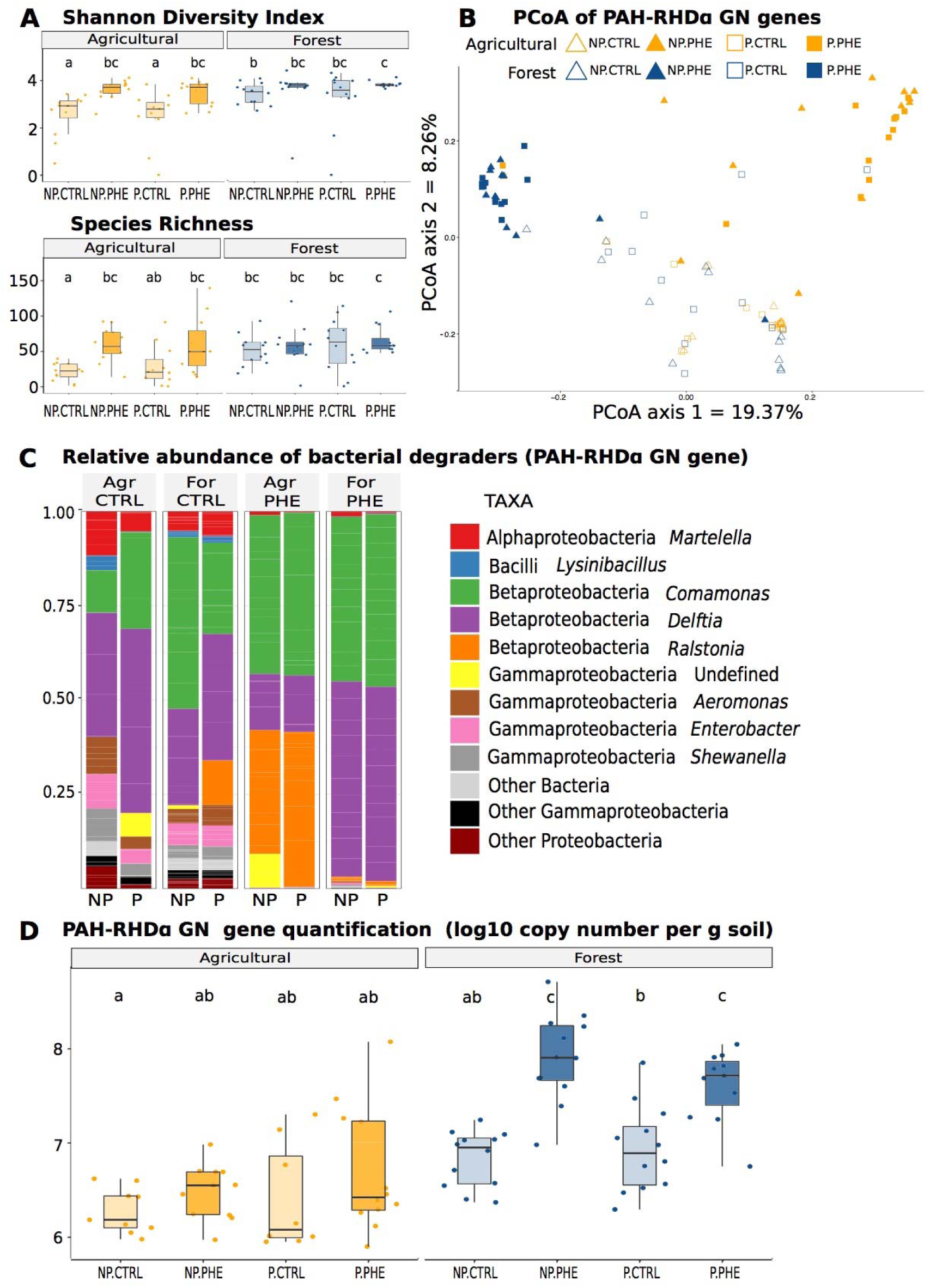
Summary of the diversity, structure and composition of PAH-degrading bacterial communities based on ASVs of the PAH-RHDα GN genes. **A**) Boxplots of Shannon diversity index and Species Richness (observed number of ASVs) by treatment. Letters denotate significant differences found with the Tukey HSD *post-hoc* test. **B**) Principal coordinates analysis (PCoA) based on Bray-Curtis dissimilarity of the relative abundance of the PAH-RHDα GN ASVs showing the effects of contamination, soil type and plant presence on the PAH-degrading Gram-negative bacterial community structure. **C**) PAH-degrading Gram-negative bacterial community composition at the class level. Only taxa with a relative abundance above 0.01 are shown. Values are averaged across treatments (n = 12). **D**) PAH-RHDα GN gene copy numbers determined by real-time PCR quantification on DNA. Legend: NP: non planted pots. P: planted pots. CTRL: non contaminated pots. PHE: pots contaminated with 50 mg kg ^-1^ of phenanthrene.

Similar trends were observed for species richness (Fig. 4a), with the number of ASVs significantly higher in the PHE agricultural soil as compared to the CTRLs, for both the P and the NP pots.

##### PAH-RHDα GN gene community structure

The PCoA of the PAH-RHDα GN gene dataset showed that, in CTRL soils, all the samples clustered more or less together, but that in the presence of contaminant, the PAH-RHDα GN communities of the two soil types clustered separately, at the two opposite ends of the first axis (Fig. 4b). PERMANOVA analysis corroborated the significant effects of soil type (R^2^ = 11.9%, p = 0.001), contamination (R^2^ = 7.2%, p value = 0.001)and the interaction of soil type and contamination (R^2^ = 8.5%, p = 0.001), and a lack of effect for the presence of plants and the related interactions (Table 1).

##### PAH-RHDα GN gene community composition

The community composition of PAH-RHDα GN genes reflected well the results of the PCoA and PERMANOVA, with similar communities across all treatments for the CTRL treatment, and largely different communities between the two soil types under PHE conditions (Fig. 4c). The PHE agricultural soils were dominated by the *Betaproteobacteria* genera *Comamonas* and *Ralstonia*, whereas in the PHE forest soils, the *Betaproteobacteria* genus *Delftia* was substituted to *Ralstonia* (Fig. 4c and Table 3).

**Table 3.** Summary of the three-way analysis of the variance (ANOVA) – with White’s correction for heteroscedasticity on the relative abundance of PAH-RHDα amplicon ASVs identified at the genus level of Gram-negative and Gram-positive bacteria. Values in bold indicate significant effects. Df, degrees of freedom. SumsOfSqs, sum of squares. MeanSqs, Mean squares. Pr(>F), p-value.

| PAH-RHDα Gram Negative |  |  |  |
| --- | --- | --- | --- |
| *Betaproteobacteria Comamonas* |  |  |  |
|  | *Df* | *F* | *P value* |
| soil | 1 | 0.9105 | 0.34314 |
| **contamination** | **1** | **6.7655** | **0.01124** |
| plant | 1 | 0.1857 | 0.66782 |
| soil:contamination | 1 | 1.3443 | 0.25006 |
| soil:plant | 1 | 0.3172 | 0.57499 |
| contamination:plant | 1 | 0.8946 | 0.34735 |
| soil:contamination:plant | 1 | 1.5475 | 0.21749 |
| Residuals | 73 |  |  |
| *Betaproteobacteria Ralstonia* |  |  |  |
|  | *Df* | *F* | *P value* |
| soil | 1 | 0.9909 | 0.32281 |
| **contamination** | **1** | **7.163** | **0.009184** |
| plant | 1 | 2.0175 | 0.159747 |
| **soil:contamination** | **1** | **25.5137** | **<0.001** |
| soil:plant | 1 | 0.2757 | 0.601121 |
| contamination:plant | 1 | 0.2601 | 0.611605 |
| soil:contamination:plant | 1 | 1.4116 | 0.238641 |
| Residuals | 73 |  |  |
| ***Betaproteobacteria Delftia*** |  |  |  |
|  | *Df* | *F* | *P value* |
| **soil** | **1** | **30.7704** | **<0.001** |
| contamination | 1 | 1.6399 | 0.204396 |
| plant | 1 | 0.016 | 0.899766 |
| **soil:contamination** | **1** | **8.4392** | **0.004857** |
| soil:plant | 1 | 0.0143 | 0.90525 |
| contamination:plant | 1 | 0.4815 | 0.489946 |
| soil:contamination:plant | 1 | 0.0542 | 0.816595 |
| Residuals | 73 |  |  |
| **PAH-RHDa Gram Positive** |  |  |  |
| ***Actinobacteria Microbacterium*** |  |  |  |
|  | *Df* | *F* | *P value* |
| **soil** | **1** | **17.863** | **<0.001** |
| contamination | 1 | 0.968 | 0.329 |
| plant | 1 | 1.155 | 0.287 |
| **soil:contamination** | **1** | **4.273** | **0.043** |
| **soil:plant** | **1** | **15.902** | **<0.001** |
| contamination:plant | 1 | 2.338 | 0.132 |
| **soil:contamination:plant** | **1** | **7.418** | **0.009** |
| Residuals | 58 |  |  |
| ***Actinobacteria Mycobacterium*** |  |  |  |
|  | *Df* | *F* | *P value* |
| soil | 1 | 1.478 | 0.229 |
| **contamination** | **1** | **43.171** | **<0.001** |
| **plant** | **1** | **7.798** | **0.007** |
| **soil:contamination** | **1** | **4.302** | **0.043** |
| soil:plant | 1 | 3.534 | 0.065 |
| contamination:plant | 1 | 0.193 | 0.662 |
| **soil:contamination:plant** | **1** | **8.304** | **0.006** |
| Residuals | 58 |  |  |

For phenanthrene concentrations, we found significant positive correlations with the relative abundance of various *Delftia* spp., (Pearsons’s correlation coefficient r ranging from 0.537 to 0.515), and significant negative correlations with the relative abundance of two *Ralstonia* spp. (Pearsons’s correlation coefficient r -0.421 and -0.426) and various *Comamonas* spp. (Pearsons’s correlation coefficient r ranging from -0.410 to -0.398) (Supp. Table 3).

##### PAH-RHDα GN gene abundance

The abundance of PAH-RHDα GN genes was significantly affected by soil type (F = 74.64, p < 0.001), contamination (F = 37.968, p < 0.001) and the interaction between these two factors (F = 11.142, p < 0.001). Both NP and P PHE forest soils harbored significantly more of PAH-RHDα GN gene copies than agricultural samples and CTRL forest samples (Fig. 4d).

#### 3.2.4. Gram-Positive bacteria degraders

##### PAH-RHDα GP gene diversity

The Shannon diversity of the PAH-RHDα GP genes was generally higher in the forest soils as compared to the agricultural soils (Fig. 5a). Kruskal-Wallis tests showed that the differences among treatments were significant (χ² = 65.466, p-value < 0.001), and the Dunn’s test revealed that the P and NP CTRL agricultural soils had a significantly lower diversity than all the other treatments, except for the P PHE agricultural soils (Fig. 5a). The trends for the abundance of ASVs observed were similar, with the P and NP CTRL agricultural soils showing a significantly lower abundance of PAH-RHDα GP ASVs compared to all the forest soil treatments.

**Figure 5.**
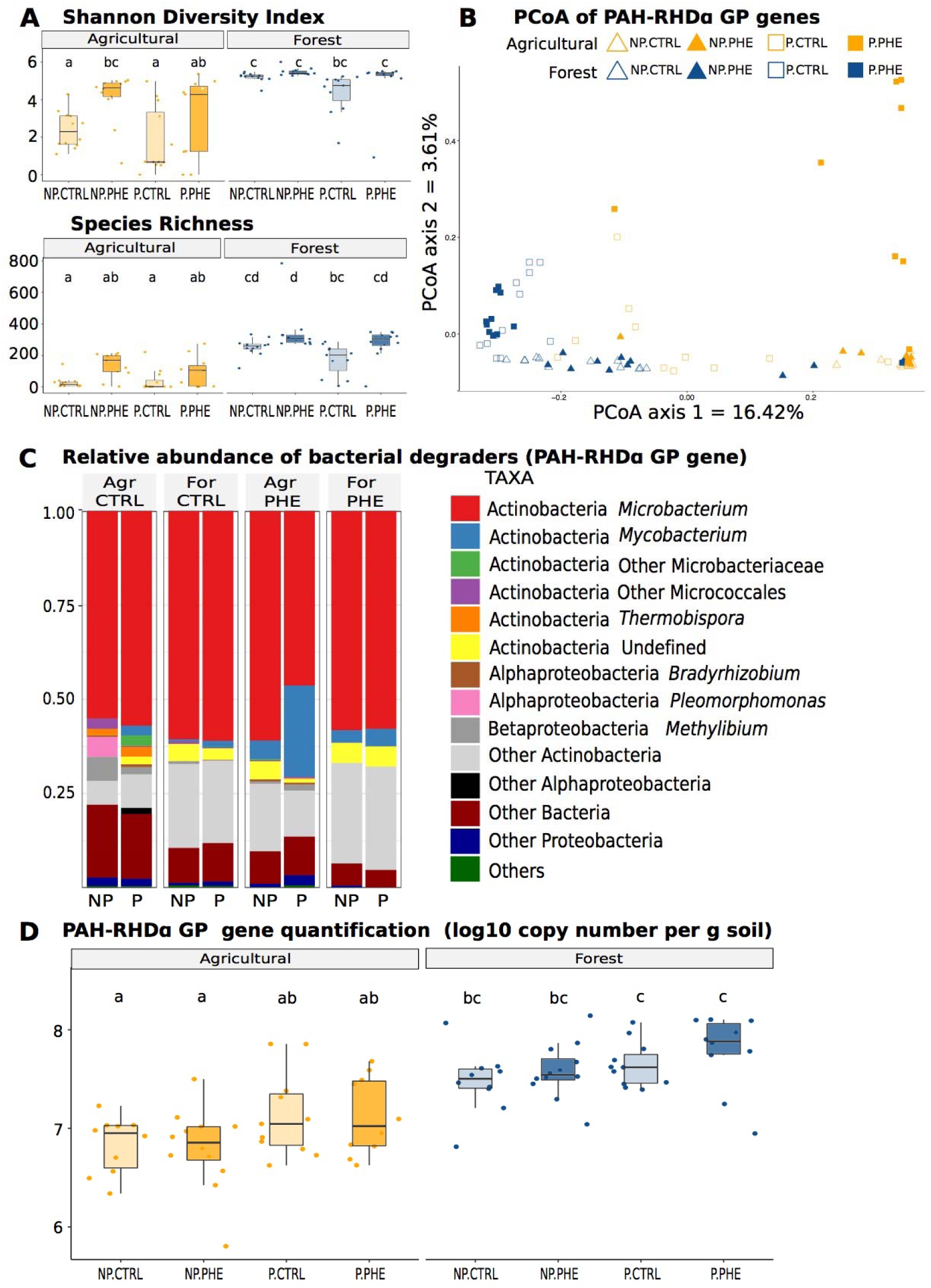
Summary of the diversity, structure and composition of PAH-degrading bacterial communities based on ASVs of the PAH-RHDα GP genes. **A**) Boxplots of Shannon diversity index and Species Richness (observed number of ASVs) by treatment. Letters denotate significant differences found with the Tukey HSD *post-hoc* test. **B**) Principal coordinates analysis (PCoA) based on Bray-Curtis dissimilarity of the relative abundance of the PAH-RHDα GP ASVs showing the effects of contamination, soil type and plant presence on the PAH-degrading Gram-positive bacterial community structure. **C**) PAH-degrading Gram-positive bacterial community composition at the class level. Only taxa with a relative abundance above 0.01 are shown. Values are averaged across treatments (n = 12). **D**) PAH-RHDα GP gene copy numbers determined by real-time PCR quantification on DNA. Legend: NP: non planted pots. P: planted pots. CTRL: non contaminated pots. PHE: pots contaminated with 50 mg kg ^-1^ of phenanthrene.

##### PAH-RHDα GP gene community structure

The PCoA ordination of the PAH-RHDα GP gene dataset showed a strong differentiation between the community structure of the two soil types (R^2^ = 14.6%, p = 0.001). The main effects of contamination (R^2^ = 3.7%, p value = 0.001) and plant presence (R^2^ = 1.9%, p = 0.019) were also significant, as well as the interaction between the three factors (R^2^ = 2.0%, p = 0.01; Table 1). For the forest treatment, there was a clear separation between the communities in NP and P pots, but not so much between the PHE and CTRL communities. In contrast, for the agricultural treatment, the PHE and CTRL communities were distinct from one another, but we also observed a separation of the NP and P communities on the first (CTRL) or second (PHE) axis of the ordination (Fig. 5b).

##### PAH-RHDα GP gene community composition

All the treatments, regardless of soil type, plant presence or contamination status were dominated by *Microbacterium* spp. ASVs (*Actinobacteria*) that accounted for more than 50% of the total amplicon sequences recovered for the PAH-RHDα GP genes (Fig. 5c). The other 50% of ASVs were divided among the phyla *Actinobacteria*, *Proteobacteria* and other unclassified Bacteria. The difference between the PHE agricultural P and NP pots observed in the PCoA could be due to the higher relative abundance of *Mycobacterium* (*Actinobacteria*) in the P PHE agricultural soils (Fig. 5c and Table 3). Also in accordance with the trends observed in the PCoA, the two different soil types showed different PAH-RHDα GP community compositions (Fig. 5c).

The top 10 most positive significant correlations with phenanthrene concentrations were for some *Microbacterium* and undefined *Actinobacteria* showing Pearsons’s correlation coefficient r ranging from 0.505 to 0.599 (Suppl. Table 3). In this case, no significant negative correlations were found.

##### PAH-RHDα GP genes quantification

There was a highly significant main effect of soil type (F = 72.674, p < 0.001), and of plant presence (F = 10.523, p = 0.002) but not of contamination on the number of PAH-RHDα GP gene copies (Fig. 5d). Forest soils showed higher numbers of PAH-RHDα GP genes than the agricultural soils, whereas the P pots harbored generally higher abundance of GP gene copies compared to the NP pots (Fig. 5d).

## 4. Discussion

Recent studies have suggested that the initial soil diversity was linked to the outcome of soil remediation and phytoremediation (Bell 2013, 2015, 2016, Yergeau 2015). Similarly, the initial soil diversity has been pointed as responsible for the variable efficiency of phytoremediation in practice (Mench et al. 2010). Based on those studies, we had hypothesized that differences in the microbiological characteristics of soil are a major constrain to PAH rhizoremediation, overriding the rhizosphere effect central to this technology. Although it is difficult to completely disentangle the effects of soil physicochemical and microbiological characteristics with our experimental design, our results confirmed that soil types had a major influence on the microbial communities and degradation functions, which resulted in different remediation patterns of phenanthrene. Indeed, soil type constrained the response of the general bacterial community to the rhizosphere effect and to contamination, with shifts mostly visible for the forest soil. Similarly, for both hydrocarbon-degradation functional genes targeted, the response of the community to contamination was constrained by soil type. These microbiological differences were mirrored in the phenanthrene degradation, which was significantly higher in the agricultural soil, with no rhizosphere effect, and lower in the forest soil, where the presence of plant significantly decreased contaminant concentrations. In contrast, fungal communities responded strongly to the rhizosphere effect, regardless of soil type and contamination, and a soil type effect was only visible in the absence of a plant.

In terms of phenanthrene degradation, the agricultural soil outperformed the forest soil. However, there were no significant differences in phenanthrene degradation in planted and unplanted pots for the agricultural soil. Both treatments efficiently degraded nearly 98% of the spiked phenanthrene at the end of the 9^th^ week, suggesting that the mere fact of distributing the soil in pots and watering them stimulated the microbial degradation enough, so that the poplar trees had no effect in agricultural soils. On the contrary, significant differences in the degradation of phenanthrene were observed between forest planted and unplanted pots. In the unplanted pots, we observed a mean total degradation around 80% as compared to 90-95% in the planted pots. This means that the effectiveness of rhizoremediation is strongly influenced by the soil type and its initial microbial diversity. Some soils appear to effectively degrade phenanthrene and potentially other PAH through the action of the microbial communities alone, without the need to introduce a plant to shift the microbial communities, whereas other soils with a different bacterial diversity and physicochemical characteristics may need stimulation from a plant in order to more effectively degrade phenanthrene. Other studies had already shown the divergences that occur during remediation of organic compounds (Bundy, Paton et Campbell 2002), with communities of microbial degraders being significantly different depending on the soil type and the original microbial diversity. When it comes specifically to PAH, the response of different soil microbial communities was also divergent, with groups of different microbial degraders arising when exposed to PAH alone or in co-contamination (Sawulski et al. 2015). These contrasting observations may result from the initial soil microbial diversity, which might explain the variable efficiency of phytoremediation across different sites. Ideally for rhizoremediation to work across sites with different soil types, the rhizosphere effect should be stronger than the soil effect. Maybe poplars do not offer this level of rhizosphere effect for PAH decontamination. Nevertheless, it is important to notice that we used a single pure contaminant in this study, which is rarely the case in non-experimental conditions. Other studies with this aa other plant species have shown the overriding effect of soil characteristics on bacterial communities in a variety of studies, as compared to plant characteristics (Bell et al. 2014; Yergeau et al. 2009; Azarbad et al. 2020). Alternatively, as fungi were shown here and elsewhere to be generally more affected by plant characteristics (Yergeau et al. 2015) and identity (Boeraeve, Honnay et Jacquemyn 2018), rhizoremediation based on fungi could be more successful across various soil types. However, metatranscriptomic studies have shown that hydrocarbon degradation genes are mainly expressed by bacteria during rhizoremediation using willows (Gonzalez 2018, Yergeau 2014, 2018).

Even though the community composition of PAH GN degraders was similar in control soils for both soil types, the communities widely differed when soils were contaminated, with *Ralstonia* and *Comamonas* dominating in the agricultural soils versus *Comamonas* and *Delftia* dominating in the forest soils. The divergence we observed in our results depending on the soil type is in agreement with the findings of other studies (Bundy, Paton et Campbell 2002; Sawulski et al. 2015). This is a good example of functional redundancy that is one of the trademark aspect of soil diversity (Dunlevy, Singleton and Aitken 2013) and can be defined as the coexistence of multiple distinct taxa or genomes capable of performing the same biochemical function (Louca et al. 2018). The relationship between biological diversity and ecological processes is far from being completely understood at a mechanistic level (Louca et al. 2018). Our study has shown that even in the context where there is an important functional redundancy among PAH degraders, the outcome at the process level, namely degradation of phenanthrene, varies widely among different soils. Similar results were previously reported, where two different soils showed specific degradation patterns for phenanthrene (Ding, Heuer et Smalla 2012). Conversely, in the unplanted pots, PAH degrader communities were similar for both soil types, but resulted in different degradation rates of phenanthrene, pointing toward an interaction between degrader communities and soil physicochemical characteristics or other microbial communities. Taken together, these results suggest that not all degraders are equal, and that depending on the pool of degraders in the original soil, of the soil physicochemical and microbiological characteristics and on the level of influence of the plant on the degraders, the outcome of the remediation will vary for similar levels of degrader diversity.

The bacterial community was mainly shaped by the soil type, but the contamination and the presence of a plant also influenced the community structure and composition. These effects were not evident at the phylum level but were clearly visible at finer taxonomical levels. For instance, *Polaromonas* and *Sphingomonas* were relatively more abundant in the PHE soils, in line with previous reports (Ding, Heuer et Smalla 2012). *Polaromonas* is a genus containing known hydrocarbon degrading species such as *P. naphthalenivorans* (Jeon et al. 2004; Ding, Heuer et Smalla 2012; Aurélie Cébron et al. 2011), and it was accordingly positively correlated with the phenanthrene concentrations in soils as it was found mainly in non-planted forest soils, where the phenanthrene degradation was probably still ongoing. Based on our qPCR quantification of PAH-RHDα GP genes, PAH degraders were significantly more abundant in planted pots as compared to unplanted pots, regardless of the contamination status of the soil. This agrees with previous studies that reported higher abundance of hydrocarbon degradation genes and transcripts in the rhizosphere of plants, even in the absence of contaminant (Cébron et al. 2011), which was suggested to be linked with the structural similarity between several plant secondary metabolites present in root exudates and PAH (Singer 2003). In contrast, the rhizosphere effect on PAH-RHDα GN genes was stronger in the contaminated soils, an effect that had been previously reported (Yergeau et al. 2014). Moreover, the soil contamination has also been found to override the willow rhizosphere effect on transcript abundance patterns in field studies (Yergeau et al. 2018).

In contrast to bacterial l and to PAH degrader communities, fungal communities were very strongly influenced by the presence of poplar and no so much by contamination. These results suggest that the fungal community is relatively stable when it comes to responding to stress caused by phenanthrene contamination, both in the rhizosphere of poplars and in the absence of plants. Most importantly, the convergence from strikingly different fungal communities in the absence of plants towards the more homogeneous communities found in rhizosphere indicate a strong plant effect. The main reason behind this trend was the complete dominance of the ectomycorrhizal fungus *Sphaerosporella* in the rhizosphere of poplars across all treatments. This fungus has been found to dominate the rhizosphere of various Salicaceae tree species in a range of pot and field experiments involving remediation of different contaminants. For instance, Dagher, Pitre and Hijri (2020) found that willow (*Salix miyabeana* clone SX67) inoculation with *S. brunnea* alone was capable of significantly improving biomass production and Ba, Zn and Cd phytoextraction with bioaccumulation in the shoot plant biomass, and an additional decrease of Cu, Pb and Sn concentrations in soil. However, Bell et al. (2015) reported that early colonization by *S. brunnea* negatively impacted the Zn accumulation efficiency of willows. In another study, the relative abundance of *S. brunnea* was positively correlated with the shoot biomass of willows (Yergeau et al. 2015). In Tardif et al (2016), *S. brunnea* showed an increase in its relative abundance in the rhizosphere of willows with increasing petroleum hydrocarbon concentrations. Similarly, willows were shown to associate more strongly with *Sphaerosporella* when growing in highly contaminated fields of Canada, but this trend was only visible for North American willows and not for Asian or European genotypes (Bell et al. 2014). Therefore, it appears that *Sphaerosporella* plays a growth promoting or stress relieving role for Salicaceae growing in contaminated soils, supporting the idea of a dual effect of the microbial community in rhizoremediation processes where some elements would play a degradative role while others would stimulate the plant growth via direct plant growth promotion, or by relieving stress (Correa-García et al. 2018). Even though this is not directly related to the degradation of contaminants, it is still of the outmost importance for rhizoremediation, as this technology is only effective where roots penetrate the soil, and an impaired growth is likely to reduce its efficiency. In the present study, since *Sphaerosporella* colonized all the rhizospheres regardless of the soil type or the contamination status, it is difficult to conclude to any positive effects on poplar growth and tolerance to contamination stress. One indication is that poplars did not present any visible stress response to phenanthrene exposition and produced equal amounts of biomass in the contaminated and non-contaminated pots. Alternatively, it could also be that the concentration of phenanthrene used was not high enough to trigger a strong stress response that would hinder plant development, as observed in previous studies (Hultgren et al 2009, Kuhn et al 2004, Ballach et al 2003). Yet again, other microbes in the rhizosphere could also have contributed by degrading enough phenanthrene in the proximity of the roots to allow for plant roots to grow without contaminant stress (Correa-García et al 2018).

In conclusion, rhizoremediation using poplars significantly improved the degradation of phenanthrene in forest soil, but not in agricultural soil. The fungal communities were the only ones strongly affected by the presence of poplar, whereas the effect of poplar on the diversity, composition, and abundance of the total bacterial and PAH degrader communities was strongly constrained by soil type, probably leading to this different degradation patterns between soils. This has important implications for rhizoremediation since these results highlight the importance of preliminary microbiological or physicochemical studies of contaminated soils in order to determine whether plant presence could improve remediation rates. Further studies could improve the predictive power of such preliminary analyses, leading to a wider adoption of this green remediation technology.

## Funding

This work was funded by the Natural Sciences and Engineering Research Council of Canada (Discovery grant RGPIN-2014-05274 and Strategic grant for projects STPGP 494702 to EY).

SCG was supported by the Research Affiliate Program from the Government of Canada. This research was enabled in part by support provided by Calcul Québec (www.calculquebec.ca) and Compute Canada.

## Credit authorship contribution statement

SCG: Conceptualization, Methodology, Field Work, Data collection, Data curation, Data Analysis, Writing and Original draft preparation. KR: Methodology, Field Work, Experimental Set up. JT: Bioinformatic analysis and Bioinformatic methods writing. AS: Conceptualization, Methodology, Experimental Set up, Supervision Reviewing and editing. EY: Conceptualization, Methodology, Supervision, Writing, Reviewing and Editing.

## Declaration of competing interest

None declared.

## Acknowledgements

The authors would like to thank the precious help provided by Denis Lachance and Gervais Pelletier during the set-up of the experiment and the maintenance of the plants both before and during the experiments, Vincenzo Corelli for developing the inhouse phenanthrene extraction methodology as well as all the members of the Laurentian Forestry Centre and the Institute Armand Frappier that contributed with ideas and discussions to improve the outcome of this study.

## Supplementary Material

**Suppl. Table 1.** Physicochemical characteristics from agricultural and forest mix soils

| Nutrient and chemical composition |  | Agricultural soil | Forest mix |
| --- | --- | --- | --- |
| pH |  | 4.6 | 5.6 |
| pH buffer |  | 5.7 | 6.0 |
| % | Organic matter | 6.1 | 6.5 |
| kg/ha | P | 31 | 81 |
|  | K | 178 | 123 |
|  | Ca | 1951 | 3338 |
|  | Mg | 130 | 134 |
| ppm | Al | 1328 | 1598 |
| ISP | P/Al | 1.0 | 2.3 |
| ppm | Mn | 20.0 | 20.0 |
|  | Cu | 1.34 | 0.83 |
|  | Zn | 3.16 | 1.96 |
|  | B | <0.1 | 0.15 |
|  | S | - | - |
|  | Fe | 280 | 301 |
| % | N total | 0.24 | 0.17 |
|  | C/N | 12.3 | 20.4 |
| ppm | N-NH<sub>4</sub> | 17.7 | 9.9 |
| ppm | N-NO<sub>3</sub> | 19.02 | 24.67 |

| Soil physical results: Granulometry |  | Agricultural soil | Forestry Mix |
| --- | --- | --- | --- |
| % | Sand | 47.6 | 87.4 |
|  | Silt | 23.0 | 5.0 |
|  | Clay | 29.5 | 7.64 |
| Texture classification |  | Clay loam | Loamy sand |
| Soil type |  | Light | Light |

**Suppl. Table 2.** Summary of the three-way analysis of the variance (ANOVA) on the relative abundances of the ITS region OTUs identified at the class level of fungi. Values in bold indicate significant effects. Df, degrees of freedom. SumsOfSqs, sum of squares. MeanSqs, Mean squares.

| ***Tremellomycetes*** | ***Df*** | ***SumOfSq*** | ***MeanOfSq*** | ***F*** | ***P value*** |
| --- | --- | --- | --- | --- | --- |
| soil | 1 | 0.011 | 0.011 | 3.673 | 0.059 |
| contamination | 1 | 0.005 | 0.005 | 1.817 | 0.181 |
| **plant** | **1** | **0.132** | **0.132** | **45.529** | **<0.001** |
| block | 1 | 0.008 | 0.008 | 2.874 | 0.094 |
| **soil:contamination** | **1** | **0.017** | **0.017** | **5.747** | **0.019** |
| soil:plant | 1 | 0.010 | 0.010 | 3.495 | 0.065 |
| contamination:plant | 1 | 0.003 | 0.003 | 0.891 | 0.348 |
| soil:contamination:plant | 1 | 0.011 | 0.011 | 3.789 | 0.055 |
| Residuals | 86 | 0.250 | 0.003 |  |  |
| ***Pezizomycetes*** | ***Df*** | ***SumOfSq*** | ***MeanOfSq*** | ***F*** | ***P value*** |
| soil | 1 | 0.000 | 0.000 | 0.021 | 0.886 |
| contamination | 1 | 0.027 | 0.027 | 2.06 | 0.155 |
| **plant** | **1** | **15.506** | **15.506** | **1186.035** | **<0.001** |
| block | 1 | 0.022 | 0.022 | 1.663 | 0.201 |
| soil:contamination | 1 | 0.023 | 0.023 | 1.753 | 0.189 |
| soil:plant | 1 | 0.001 | 0.001 | 0.055 | 0.816 |
| contamination:plant | 1 | 0.012 | 0.012 | 0.954 | 0.331 |
| soil:contamination:plant | 1 | 0.017 | 0.017 | 1.312 | 0.255 |
| Residuals | 86 | 1.124 | 0.013 |  |  |
| ***Mortierellomycotina*** | ***Df*** | ***SumOfSq*** | ***MeanOfSq*** | ***F*** | ***P value*** |
| **soil** | **1** | **0.504** | **0.504** | **48.379** | **<0.001** |
| contamination | 1 | 0.001 | 0.001 | 0.053 | 0.818 |
| **plant** | **1** | **1.373** | **1.373** | **131.763** | **<0.001** |
| block | 1 | 0.017 | 0.017 | 1.599 | 0.210 |
| **soil:contamination** | **1** | **0.049** | **0.049** | **4.697** | **0.033** |
| **soil:plant** | **1** | **0.287** | **0.287** | **27.518** | **<0.001** |
| contamination:plant | 1 | 0.000 | 0.000 | 0.036 | 0.851 |
| soil:contamination:plant | 1 | 0.059 | 0.059 | 5.702 | 0.019 |
| Residuals | 86 | 0.896 | 0.010 |  |  |
| ***Leotiomyces*** | ***Df*** | ***SumOfSq*** | ***MeanOfSq*** | ***F*** | ***P value*** |
| **soil** | **1** | **0.090** | **0.090** | **5.194** | **0.025** |
| contamination | 1 | 0.002 | 0.002 | 0.125 | 0.724 |
| **plant** | **1** | **0.450** | **0.450** | **26.133** | **<0.001** |
| block | 1 | 0.002 | 0.002 | 0.141 | 0.708 |
| soil:contamination | 1 | 0.017 | 0.017 | 0.956 | 0.331 |
| soil:plant | 1 | 0.006 | 0.006 | 0.335 | 0.564 |
| contamination:plant | 1 | 0.024 | 0.024 | 1.409 | 0.238 |
| soil:contamination:plant | 1 | 0.018 | 0.018 | 1.048 | 0.309 |
| Residuals | 86 | 1.482 | 0.017 |  |  |
| ***Agaricomycetes*** | ***Df*** | ***SumOfSq*** | ***MeanOfSq*** | ***F*** | ***P value*** |
| **soil** | **1** | **0.074** | **0.074** | **34.734** | **<0.001** |
| contamination | 1 | 0.001 | 0.001 | 0.258 | 0.613 |
| **plant** | **1** | **0.146** | **0.146** | **68.675** | **<0.001** |
| block | 1 | 0.000 | 0.000 | 0.047 | 0.829 |
| soil:contamination | 1 | 0.000 | 0.000 | 0.025 | 0.875 |
| **soil:plant** | **1** | **0.032** | **0.032** | **15.072** | **<0.001** |
| contamination:plant | 1 | 0.001 | 0.001 | 0.313 | 0.578 |
| soil:contamination:plant | 1 | 0.000 | 0.000 | 0.053 | 0.819 |
| Residuals | 86 | 0.182 | 0.002 |  |  |

**Suppl. Table 3.**
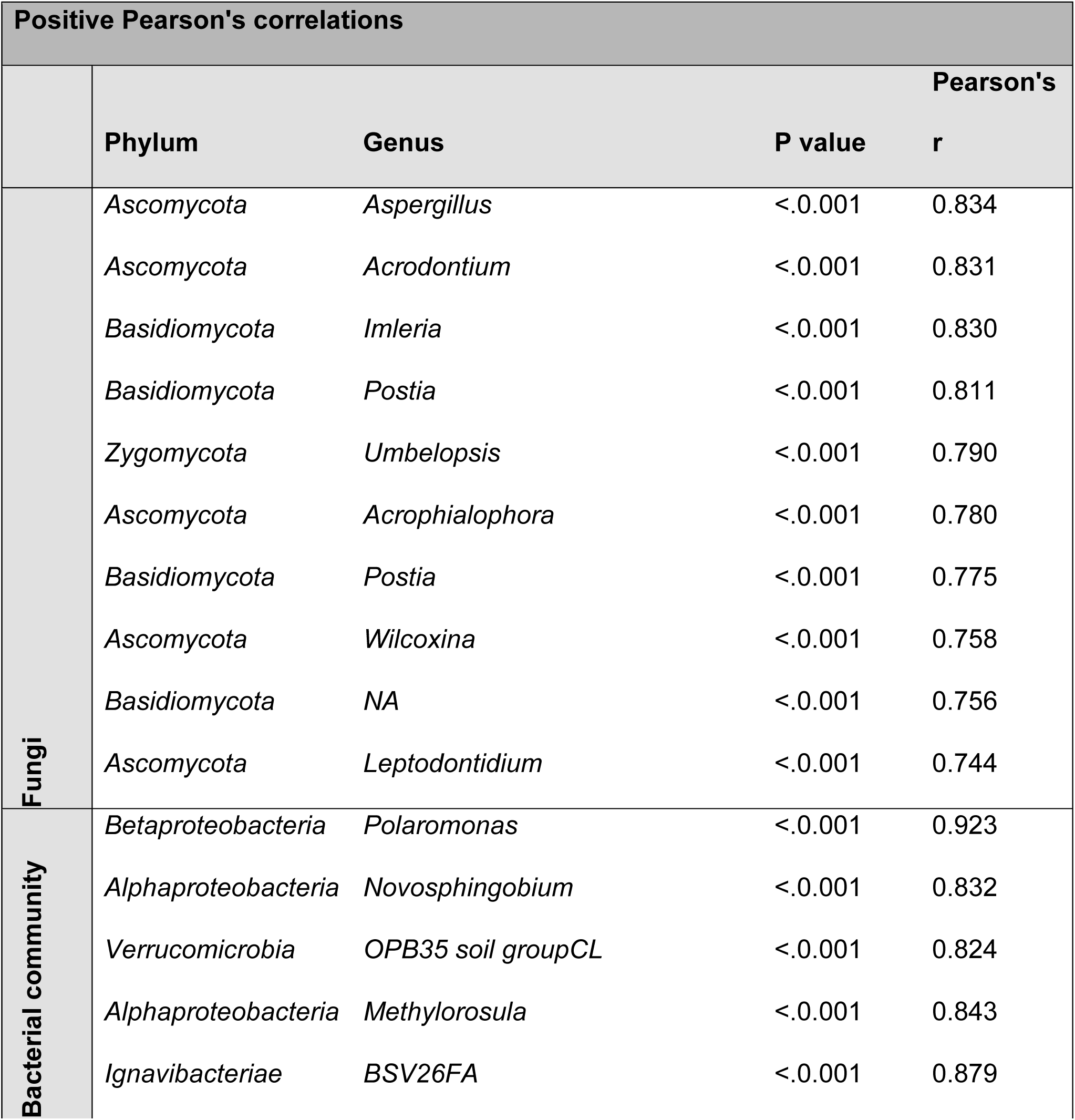
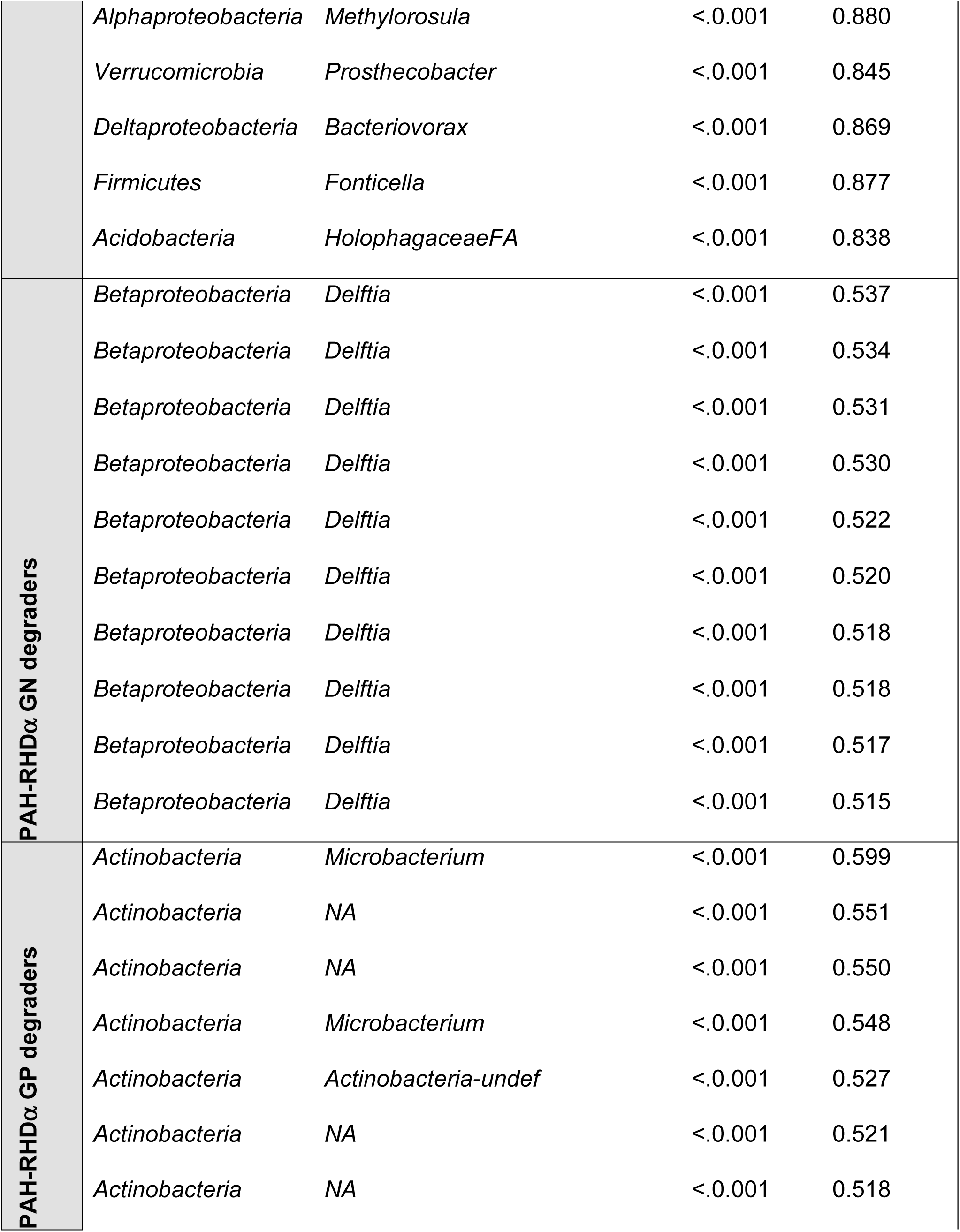
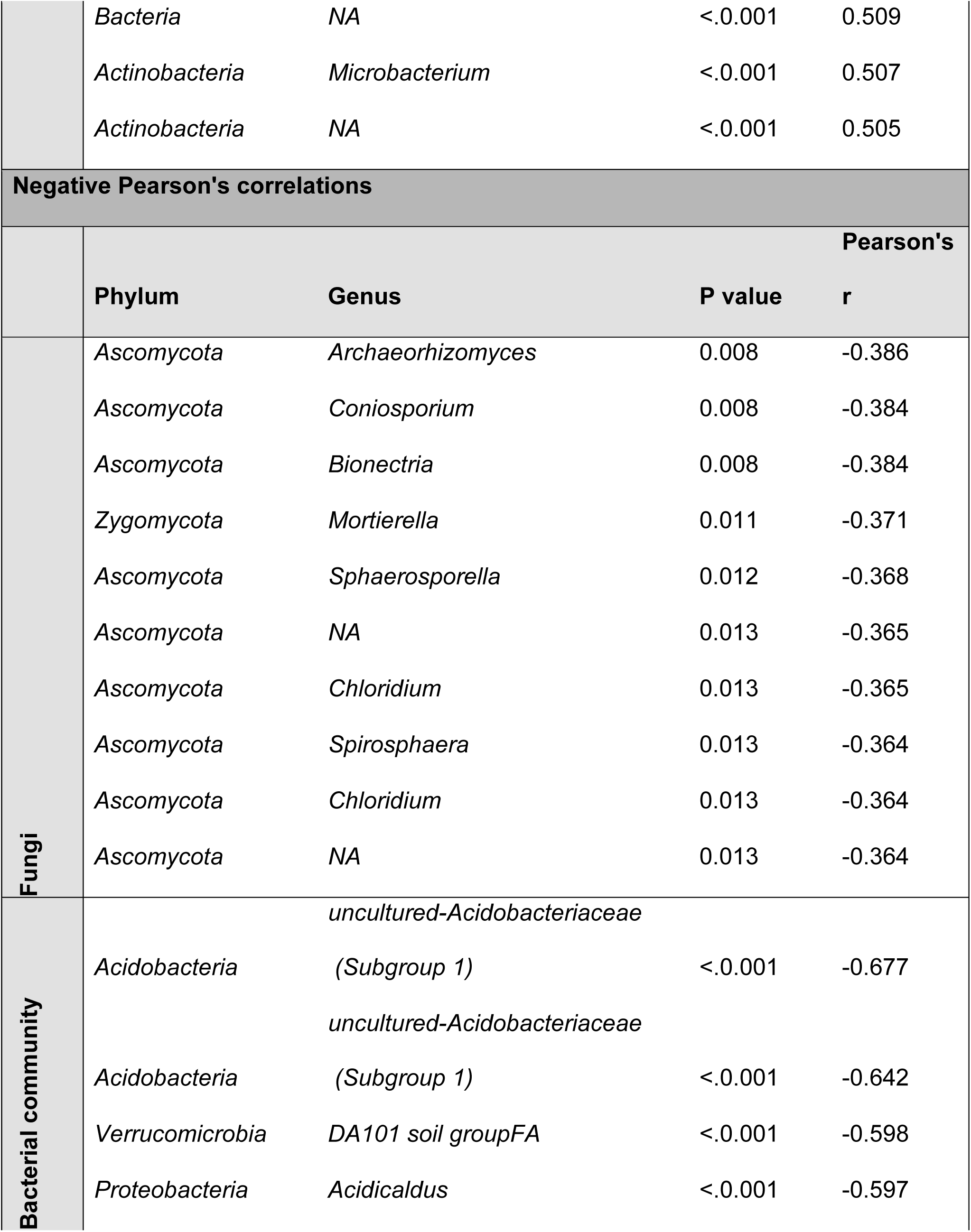
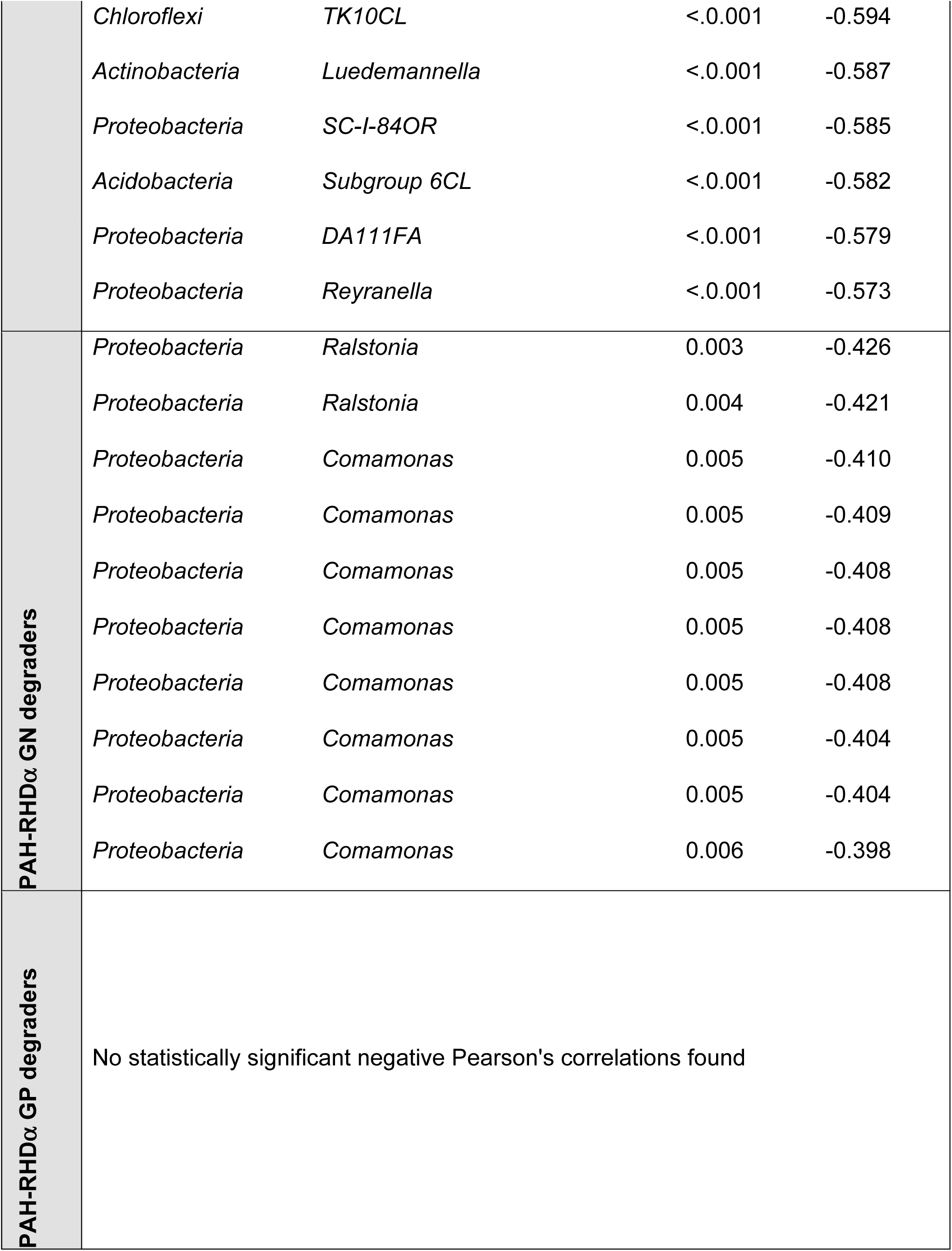
Statistically significant Pearson’s coefficients of correlation r between the relative abundance of bacterial and fungal OTUs or PAH bacterial degraders ASVs identified at the genus level and the quantity of phenanthrene (mg kg ^-1^) found in soil.

**Suppl. Table 4.** Summary of the three-way analysis of the variance (ANOVA) on the relative abundances of the 16S rRNA gene OTUs identified at the phylum level of bacteria. Values in bold indicate significant effects. Df, degrees of freedom. SumsOfSqs, sum of squares. MeanSqs, Mean squares.

| *Acidobacteria* | Df | SumOfSq | MeanOfSq | F | P value |
| --- | --- | --- | --- | --- | --- |
| **soil** | **1** | **0.15681** | **0.15681** | **59.017** | **<0.001** |
| **contamination** | 1 | 0.00299 | 0.00299 | 1.127 | 0.291 |
| **plant** | 1 | 0.00024 | 0.00024 | 0.090 | 0.765 |
| **block** | 1 | 0.00044 | 0.00044 | 0.165 | 0.686 |
| **soil:contamination** | 1 | 0.00312 | 0.00312 | 1.174 | 0.282 |
| **soil:plant** | **1** | **0.02612** | **0.02612** | **9.830** | **0.002** |
| **contamination:plant** | 1 | 0.00018 | 0.00018 | 0.067 | 0.796 |
| **soil:contamination:plant** | 1 | 0.00015 | 0.00015 | 0.056 | 0.813 |
| **Residuals** | 87 | 0.23117 | 0.00266 |  |  |
| *Actinobacteria* | Df | SumOfSq | MeanOfSq | F | P value |
| **soil** | 1 | 0.00060 | 0.00060 | 0.606 | 0.438 |
| **contamination** | 1 | 0.00106 | 0.00106 | 1.065 | 0.305 |
| **plant** | **1** | **0.02038** | **0.02038** | **20.577** | **<0.001** |
| **block** | 1 | 0.00279 | 0.00279 | 2.817 | 0.097 |
| **soil:contamination** | 1 | 0.00037 | 0.00037 | 0.371 | 0.544 |
| **soil:plant** | **1** | **0.00350** | **0.00350** | **3.534** | **0.064** |
| **contamination:plant** | 1 | 0.00014 | 0.00014 | 0.138 | 0.711 |
| **soil:contamination:plant** | 1 | 0.00012 | 0.00012 | 0.118 | 0.732 |
| **Residuals** | 87 | 0.08618 | 0.00099 |  |  |
| ***Alphaproteobacteria*** | **Df** | **SumOfSq** | **MeanOfSq** | **F** | **P value** |
| **soil** | **1** | **0.02187** | **0.02187** | **32.006** | **<0.001** |
| **contamination** | 1 | 0.00162 | 0.00162 | 2.374 | 0.127 |
| **plant** | **1** | **0.00416** | **0.00416** | **6.092** | **0.016** |
| **block** | 1 | 0.00050 | 0.00050 | 0.734 | 0.394 |
| **soil:contamination** | **1** | **0.00404** | **0.00404** | **5.911** | **0.017** |
| **soil:plant** | **1** | **0.00721** | **0.00721** | **10.557** | **0.002** |
| **contamination:plant** | 1 | 0.00002 | 0.00002 | 0.026 | 0.873 |
| **soil:contamination:plant** | 1 | 0.00108 | 0.00108 | 1.577 | 0.213 |
| **Residuals** | 87 | 0.05945 | 0.00068 |  |  |
| ***Bacteroidetes*** | **Df** | **SumOfSq** | **MeanOfSq** | **F** | **P value** |
| **soil** | **1** | **0.10442** | **0.10442** | **275.678** | **<0.001** |
| **contamination** | **1** | **0.00137** | **0.00137** | **3.607** | **0.061** |
| **plant** | **1** | **0.00471** | **0.00471** | **12.433** | **0.001** |
| **block** | **1** | **0.00232** | **0.00232** | **6.119** | **0.015** |
| **soil:contamination** | 1 | 0.00007 | 0.00007 | 0.184 | 0.669 |
| **soil:plant** | **1** | **0.00663** | **0.00663** | **17.503** | **<0.001** |
| **contamination:plant** | 1 | 0.00000 | 0.00000 | 0.009 | 0.926 |
| **soil:contamination:plant** | 1 | 0.00025 | 0.00025 | 0.654 | 0.421 |
| **Residuals** | 87 | 0.03295 | 0.00038 |  |  |

| ***Betaproteobacteria*** | **Df** | **SumOfSq** | **MeanOfSq** | **F** | **P value** |
| --- | --- | --- | --- | --- | --- |
| **soil** | **1** | **0.23470** | **0.23467** | **41.812** | **<0.001** |
| **contamination** | 1 | 0.00010 | 0.00006 | 0.010 | 0.919 |
| **plant** | **1** | **0.03190** | **0.03187** | **5.678** | **0.019** |
| **block** | 1 | 0.00290 | 0.00292 | 0.521 | 0.472 |
| **soil:contamination** | 1 | 0.00010 | 0.00010 | 0.019 | 0.892 |
| **soil:plant** | **1** | **0.12150** | **0.12150** | **21.648** | **<0.001** |
| **contamination:plant** | 1 | 0.00170 | 0.00168 | 0.300 | 0.585 |
| **soil:contamination:plant** | 1 | 0.00850 | 0.00846 | 1.508 | 0.223 |
| **Residuals** | 87 | 0.48830 | 0.00561 |  |  |
| ***Chloroflexi*** | **Df** | **SumOfSq** | **MeanOfSq** | **F** | **P value** |
| **soil** | **1** | **0.25941** | **0.25941** | **548.362** | **<0.001** |
| **contamination** | 1 | 0.00097 | 0.00097 | 2.052 | 0.156 |
| **plant** | **1** | **0.00152** | **0.00152** | **3.211** | **0.077** |
| **block** | 1 | 0.00029 | 0.00029 | 0.604 | 0.439 |
| **soil:contamination** | 1 | 0.00011 | 0.00011 | 0.237 | 0.627 |
| **soil:plant** | 1 | 0.00030 | 0.00030 | 0.632 | 0.429 |
| **contamination:plant** | 1 | 0.00000 | 0.00000 | 0.009 | 0.923 |
| **soil:contamination:plant** | 1 | 0.00006 | 0.00006 | 0.120 | 0.730 |
| **Residuals** | 87 | 0.04116 | 0.00047 |  |  |
| ***Deltaproteobacteria*** | **Df** | **SumOfSq** | **MeanOfSq** | **F** | **P value** |
| **soil** | **1** | **0.00529** | **0.00529** | **42.913** | **<0.001** |
| **contamination** | 1 | 0.00009 | 0.00009 | 0.717 | 0.399 |
| **plant** | 1 | 0.00014 | 0.00014 | 1.148 | 0.287 |
| **block** | 1 | 0.00036 | 0.00036 | 2.930 | 0.091 |
| **soil:contamination** | 1 | 0.00014 | 0.00014 | 1.164 | 0.284 |
| **soil:plant** | 1 | 0.00020 | 0.00020 | 1.617 | 0.207 |
| **contamination:plant** | 1 | 0.00004 | 0.00004 | 0.287 | 0.594 |
| **soil:contamination:plant** | 1 | 0.00003 | 0.00003 | 0.247 | 0.620 |
| **Residuals** | 87 | 0.01072 | 0.00012 |  |  |
| ***Firmicutes*** | **Df** | **SumOfSq** | **MeanOfSq** | **F** | **P value** |
| **soil** | **1** | **0.0026** | **0.0026** | **28.414** | **<0.001** |
| **contamination** | **1** | **0.0006** | **0.0006** | **6.677** | **0.011** |
| **plant** | 1 | 0.0002 | 0.0002 | 2.403 | 0.125 |
| **block** | 1 | 0.0000 | 0.0000 | 0.007 | 0.934 |
| **soil:contamination** | **1** | **0.0005** | **0.0005** | **5.148** | **0.026** |
| **soil:plant** | 1 | 0.0001 | 0.0001 | 0.936 | 0.336 |
| **contamination:plant** | **1** | **0.0003** | **0.0003** | **3.260** | **0.075** |
| **soil:contamination:plant** | 1 | 0.0000 | 0.0000 | 0.445 | 0.506 |
| **Residuals** | 87 | 0.0080 | 0.0001 |  |  |
| ***Gammaproteobacteria*** | **Df** | **SumOfSq** | **MeanOfSq** | **F** | **P value** |
| **soil** | **1** | **0.0152** | **0.0152** | **73.862** | **<0.001** |
| **contamination** | **1** | **0.0011** | **0.0011** | **5.283** | **0.024** |
| **plant** | 1 | 0.0000 | 0.0000 | 0.053 | 0.819 |
| **block** | 1 | 0.0000 | 0.0000 | 0.191 | 0.663 |
| **soil:contamination** | **1** | **0.0008** | **0.0008** | **3.918** | **0.051** |
| **soil:plant** | 1 | 0.0000 | 0.0000 | 0.059 | 0.808 |
| **contamination:plant** | 1 | 0.0004 | 0.0004 | 1.809 | 0.182 |
| **soil:contamination:plant** | 1 | 0.0000 | 0.0000 | 0.060 | 0.806 |
| **Residuals** | 87 | 0.0179 | 0.0002 |  |  |
| ***Gemmatimonadetes*** | **Df** | **SumOfSq** | **MeanOfSq** | **F** | **P value** |
| **soil** | **1** | **0.0024** | **0.0024** | **29.953** | **<0.001** |
| **contamination** | **1** | **0.0011** | **0.0011** | **14.454** | **<0.001** |
| **plant** | 1 | 0.0000 | 0.0000 | 0.140 | 0.709 |
| **block** | 1 | 0.0000 | 0.0000 | 0.184 | 0.669 |
| **soil:contamination** | 1 | 0.0000 | 0.0000 | 0.222 | 0.639 |
| **soil:plant** | 1 | 0.0002 | 0.0002 | 2.701 | 0.104 |
| **contamination:plant** | 1 | 0.0000 | 0.0000 | 0.072 | 0.789 |
| **soil:contamination:plant** | 1 | 0.0003 | 0.0003 | 3.239 | 0.075 |
| **Residuals** | 87 | 0.0069 | 0.0001 |  |  |
| ***Planctomycetes*** | **Df** | **SumOfSq** | **MeanOfSq** | **F** | **P value** |
| **soil** | **1** | **0.0003** | **0.0003** | **5.550** | **0.021** |
| **contamination** | **1** | **0.0003** | **0.0003** | **5.978** | **0.017** |
| **plant** | **1** | **0.0004** | **0.0004** | **8.910** | **0.004** |
| **block** | 1 | 0.0000 | 0.0000 | 0.039 | 0.844 |
| **soil:contamination** | 1 | 0.0001 | 0.0001 | 2.353 | 0.129 |
| **soil:plant** | **1** | **0.0003** | **0.0003** | **7.238** | **0.009** |
| **contamination:plant** | 1 | 0.0000 | 0.0000 | 0.465 | 0.497 |
| **soil:contamination:plant** | 1 | 0.0001 | 0.0001 | 1.937 | 0.168 |
| **Residuals** | 87 | 0.0041 | 0.0000 |  |  |
| **Others** | **Df** | **SumOfSq** | **MeanOfSq** | **F** | **P value** |
| **soil** | **1** | **0.0014** | **0.0014** | **36.033** | **<0.001** |
| **contamination** | 1 | 0.0000 | 0.0000 | 1.214 | 0.274 |
| **plant** | 1 | 0.0000 | 0.0000 | 0.073 | 0.787 |
| **block** | 1 | 0.0000 | 0.0000 | 0.322 | 0.572 |
| **soil:contamination** | 1 | 0.0001 | 0.0001 | 1.252 | 0.266 |
| **soil:plant** | 1 | 0.0000 | 0.0000 | 0.166 | 0.685 |
| **contamination:plant** | 1 | 0.0000 | 0.0000 | 0.712 | 0.401 |
| **soil:contamination:plant** | 1 | 0.0001 | 0.0001 | 2.251 | 0.137 |
| **Residuals** | 87 | 0.0035 | 0.0000 |  |  |

**Suppl. Table 5.** Read counts reports por whole bacterial and fungi communities (OTUs) and for PAH Gram-negative and Gram-positive bacterial communities (ASVs).

|  | 16S rRNA | ITS region | PAH-RHD GN | PAH-RHD GP |
| --- | --- | --- | --- | --- |
| total_reads | 18,877,788 | 16,168,948 | 10,559,056 | 7,055,496 |
| contaminants_reads | 11,066 | 557,756 | 631,164 | 2,961,494 |
| phix_reads | 2,136 | 892 | 1,458 | 2,872 |
| non_contam_non_phix_reads | 18,864,586 | 15,610,300 | 9,926,434 | 4,091,130 |
| non_contam_non_phix_reads_1 | 9,432,293 | 7,805,150 | 4,963,217 | 2,045,565 |
| non_contam_non_phix_reads_2 | 9,432,293 | 7,805,150 | 4,121,810 | 819,318 |
| assembled_reads | 9,305,398 | 5,981,093 |  |  |
| assembled_reads_QC_passed | 8,535,756 | 5,529,429 |  |  |
| Clustered | 4,646,772 | 5,245,486 | 3,543,681 | 538,575 |
|  | sequences in | sequences in | sequences in | sequences in |
|  | 21,182 | 3,464 | 1,438 | 3,019 clusters |
|  | clusters | clusters | clusters |  |
| Classified clusters | 4,646,766 | 5,240,163 | 3,543,681 | 538,513 |
|  | sequences | sequences | sequences | sequences |
|  | packed in | packed in | packed in | packed in |
|  | 21,181 cluster | 3,327 cluster | 1,438 cluster | 3,017 cluster |

